# Abrogation and Homeostatic Restoration of IgE Responses by a Universal IgE Allergy CTL Vaccine—The Three Signal Self/Non-Self/Self (S/NS/S) Model

**DOI:** 10.1101/2023.10.12.561777

**Authors:** Swey-Shen Chen, Hailan Zhang

**Author notes:** Swey-Shen Chen, Lead and Communicating Author.

## Abstract

Natural IgE cytotoxic peptides (nECPs), which are derived from the constant domain of the heavy chain of human IgE producing B cells via endoplasmic reticulum (ER) stress, are decorated onto MHC class 1a molecules (MHCIa) as unique biomarkers for CTL (cytotoxic T lymphocyte)-mediated immune surveillance. Human IgE exhibits only one isotype and lacks polymorphisms; IgE is pivotal in mediating diverse, allergen-specific allergies. Therefore, by disrupting self-IgE tolerance via costimulation, the cytotoxic T lymphocytes (CTLs) induced by nECPs can serve as universal allergy vaccines (UAVs) in humans to dampen IgE production mediated by diverse allergen-specific IgE-secreting B cells and plasma cells expressing surface nECP-MHCIa as targets. The study herein has enabled the identification of nECPs produced through the correspondence principle ^1, 2^. Furthermore, nECP-tetramer-specific CTLs were found to be converted into CD4 Tregs that restored IgE competence via the homeostatic principle, mediated by SREBP-1c suppressed DCs. Thus, nECPs showed causal efficacy and safety as UAVs for treating type I hypersensitivity IgE-mediated allergies. The applied vaccination concept presented provides the foundation to unify, integrate through a singular class of tetramer-specific TCR clonotypes. The three signal model is proposed on the mechanisms underlying central tolerance, breaking tolerance and regaining peripheral tolerance via homeostasis concerning nECP as an efficacious and safe UAV to treat type I IgE-mediated hypersensitivity.

**One Sentence Summary:** Human IgE self-peptides are identified as universal allergy vaccines that inhibit IgE synthesis while allowing homeostatic IgE recovery.

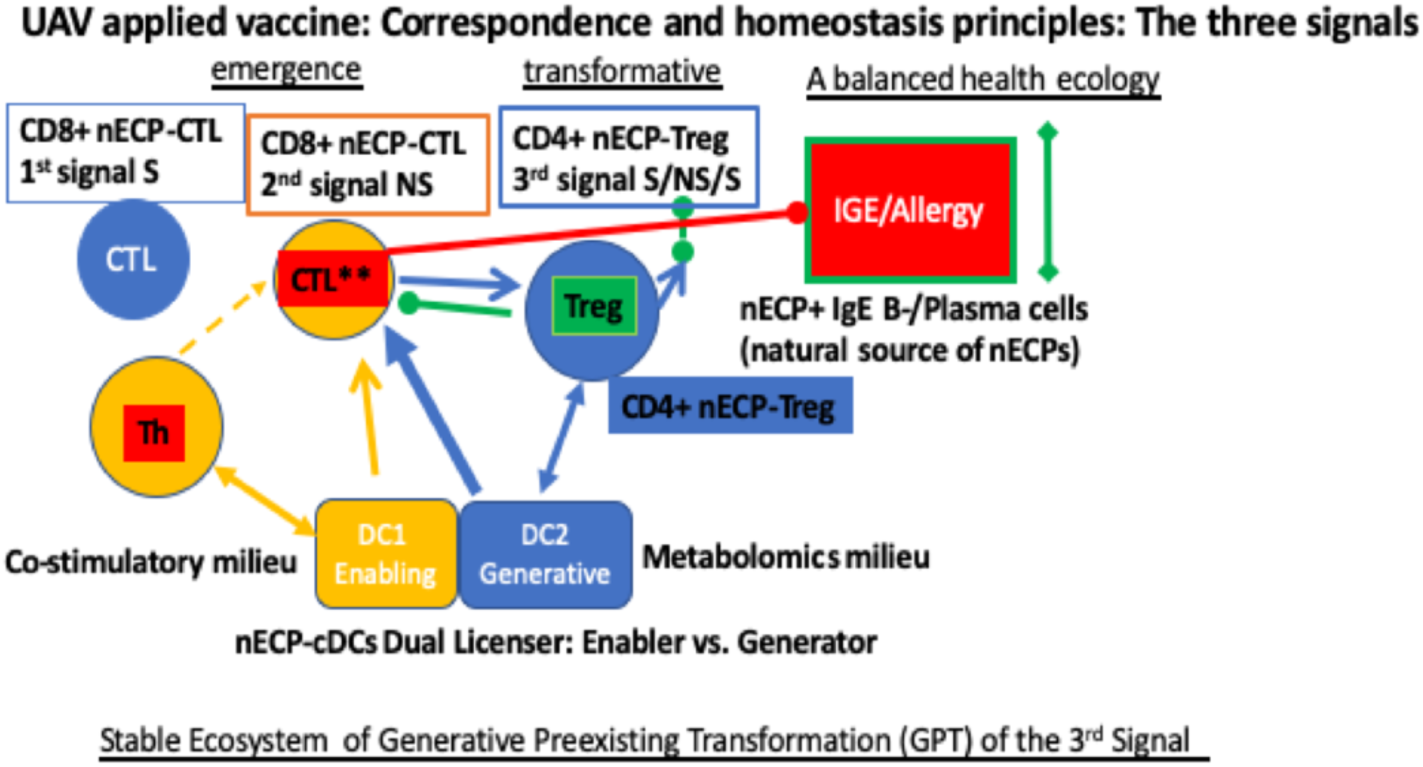

Graphic abstract text

Three cell S/NS/S model of Universal Allergy Vaccines (UAV): Natural IgE peptides (nECPs) presented by enabler DCs break central IgE tolerance (Self), leading to CTLs that inhibit IgE production (Non-self). Generative DCs converted by the metabolic milieu transform the pre-existing nECP-specific CTLs into nECP-specific Tregs leading to homeostatic recovery of IgE competence (S).

## Introduction

IgE is a critical class of immunoglobulin playing a crucial gatekeeper function to amplify IgE-mediated defense to parasites and other functions in the mucosal respiratory and the GI tract microenvironment. On the other hand, dysregulated IgE production is the central culprit for mediating type 2 immediate hypersensitivity, accounting for allergic asthma, rhinitis, urticaria, peanut allergy and food allergy in more than 40% of the worldwide population.

During the perinatal period in rodents, IgE system, not other isotypes, is highly autoreactive, and pre-mature exposure to IgE or externally in soluble form without adjuvant or conjugated on splenic APCs in particular to newborn to up to 72 h perinates lead to anti-IgE, and suppressive cell-mediated immunity that result in life-long lack of allergen-specific IgE production and profound life-long total IgE deficiency, while natural tolerance to IgE is completed within one week after birth concordant with the appearance of endogenous IgE in the laboratory ^1, 3-7^. Dampening the levels of IgE in allergen-sensitized patients regardless of allergen specificity is considered the goal of a universal allergy therapy.

It has been shown recently that the generation of chimeric antigen receptor (CAR) T cells using antibodies directed against the proteins produced by cells that cause fibrosis can attenuate fibrosis in multiple organs ^8^. However, the anti-IgE antibody Xolair, which targets FcεRIα receptor-sequestered determinants, does not recognize surface IgE on IgE-committed B cells and plasma cells through an antibody-dependent cellular cytotoxicity (ADCC) mechanism ^9^. Therefore, we set out to identify naturally produced IgE cytotoxic peptide biomarkers that are associated with MHCIa supertypes and can induce CTLs to inhibit IgE production by IgE B cell lineages decorated with self-IgE markers as well as to ensure homeostatic IgE recovery via natural IgE cytotoxic peptides (nECP)-based vaccination. Rodent and human exhibit one single IgE isotype without polymorphism. Any vaccine direct against the unique isotype-specific B cell or T cell epitope offer the promise of a universal IgE or allergy vaccine (UAV). The concept of using unique IgE self-biomarkers on the IgE-producing B cells and plasma cells as CTL targets was first validated in rodents. IgE production was inhibited by CTLs in adult vaccinated with bone marrow-derived dendritic cells (BMDCs) co-presenting rodent natural IgE peptides, peptide I ^7^ restricted to K^d^/D^b^ along with a conventional helper peptide in the laboratory ^10, 11^.

The vaccines are focused on HLA-A 2.01 and HLA-B 7.02 MHCIa supertypes that are prevalent among approximately 48% of the human population ^12^. Herein we characterize in the laboratory three nonameric nECPs, A32 for HLA-A2.01 and SP-1 and SP-2 for HLA-B 7.02, and coupled with helper peptides in breaking tolerance, which induce nECP tetramer-specific CD8 CTLs, via the correspondence principle, e.g., the vaccine epitopes displayed on MoDCs also display on IgE producing cells. Thus, nECP- specific CTLs targeting human IgE producing cells that profoundly inhibit human IgE production by IgE B cell lineages in human peripheral blood mononuclear cells (PBMCs) *in vitro.* Moreover, the correspondence principle is ascertained in chimeric human IgE production in triple transgenic animals, expressing human IgE-producing on B cells, expressing nECP presented by HLA-A2.01 and HLA-B7.02, and nECP-specific HLA-A2.01 and HLA-B7.02 that suppressing chimeric human IgE production in triple transgenic rodents. Furthermore, we showed that via the homeostatic regulatory principle, nECP-tetramer-specific CTLs can be transformed into nECP-tetramer-specific CD4 Tregs, which decrease the levels of CD8 CTLs and spare human IgE B cells, thus restoring IgE production. Based on these two vaccine principles and observations in the laboratory, we propose that a novel class of nECPs can be used to generate universal allergy vaccines (UAVs; nECPs-UAVs) that have strong causal efficacy in decreasing human IgE production to treat IgE-mediated hypersensitivity diseases. The applied vaccination concept presented provides the foundation to unify, integrate through a singular class of tetramer-specific TCR clonotypes. The three signal model is proposed on the mechanisms underlying central tolerance, breaking tolerance and regaining peripheral tolerance via homeostasis concerning nECP as an efficacious and safe UAV to treat type I IgE-mediated hypersensitivity. The self-biomarkers as targets for self-reactive CTLs is considered may also be relevant for treating allergies, autoimmunity, and cancers ^13^.

## Results

### 1. The principle of correspondence of nECP tetramer-specific CTLs that can target IgE-producing cells

PBMCs from multiple human donors were typed by primer sequencing of HLA-B7.02 and/or HLA-A2.01, which were comprehensively screened using the sequences of hundreds of synthetic peptides (IEBD/MHCPan4.0/BIMAS programs) for binding and/or CTL induction using promiscuous helper T-cell peptides. SP-1 and SP-2 human IgE peptides strongly increase by one log of shift of fluorescent intensity of HLA-B7.02 on HLA-B7.02-transfected TAP-1/2-deficient T2 cells, kindly provided by Dr. Lutz at University of Kentucky ^14^ comparable to the positive control EBV peptide, whereas SP- 4,-5, 6, 11 did not increase HLA-B7.02 surface expression. The SP4 control peptides were repeated twice to show the precision of the negative baseline in contrast to the log shift of the highly efficient SP-1 and SP-2 to upregulate HLA-B7.02 ^14^ (Fig. 1A) (Suppl. Extended Data Table 1). Since the human A32 IgE peptide did not upregulate surface MHCIa upregulation due to low affinity binding, we resorted a direct T-cell activation assay: a single candidate A32 was selected following a large array of putative HLA- A2.01 binding synthetic peptides. Herein as shown in Fig. 1B, the A32 peptide (HLA- A2.01), presented by endogenous FTL-3-stimulated classic dendritic cells (cDCs) according to methods of V. Appay ^15, 16^ (and later by monocyte-derived dendritic cells (MoDCs), not shown) ^17^ together with the promiscuous helper T-cell peptide, PADRE ^18^ was shown to more potently induce the production of IFN-γ/CD137 (4-1BB) + CD8 T cells ^19^ (095% double positive, 1.19% CD137+, and 3.81% IFN-γ+, total 5.95%) more potent than the canonical positive control melanoma ELA-10 peptide ^20^ for HLA-A2.01 (2.94%), despites its low affinity unable to upregulate surface HLA-A2.01 in TAP-1/2 deficient T2, nor repeated failed attempts for tetramer formation with HLA-A2.01 (Fig. 1B) (Suppl. Extended Data: The three-step method) ^16^. It is crucial to monitor nECP- specific CD8 T cells levels for the vaccination purpose. Thus the high affinity HLA-B7.02 biding SP-1 and SP-2 PE-conjugated tetramers were prepared (provided by the NIH tetramer facility, Atlanta, GA). Indeed, we detected a surprisingly high frequency of major clonotypes (11.6%) of tetramer-positive CD8 T cells expressing CD137 in the FLT-3-stimulated PBMCs as previously used for A32 activated clonotypes at 5.9% (Fig. 1B). Moreover, Fig. 1C showed that SP-1, and SP-2 nECP activated TCR clonotypes increased approximately 3.8-fold (44.6%) after treatment with the AMPK activator AICAR/metformin, glucose and lipid sensors (AMPKm) ^21 12323, 22^. Notably, significant numbers of tetramer-gated cells were double positive for CD137 and IFN-γ expression (7.24%), which increased approximately 3.3-fold (24%) in the presence of AMPK activators (Fig. 1D).

**Fig. 1.**
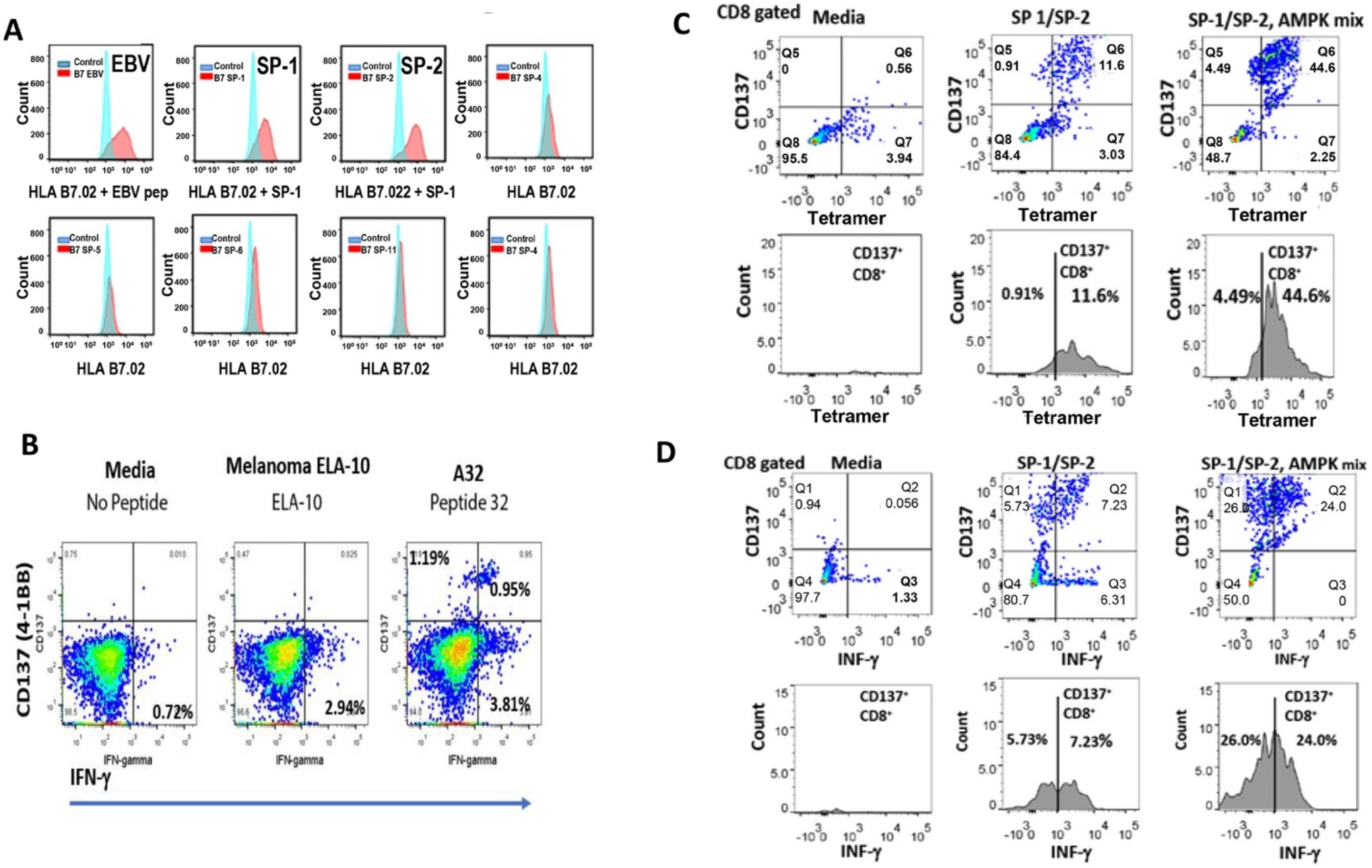
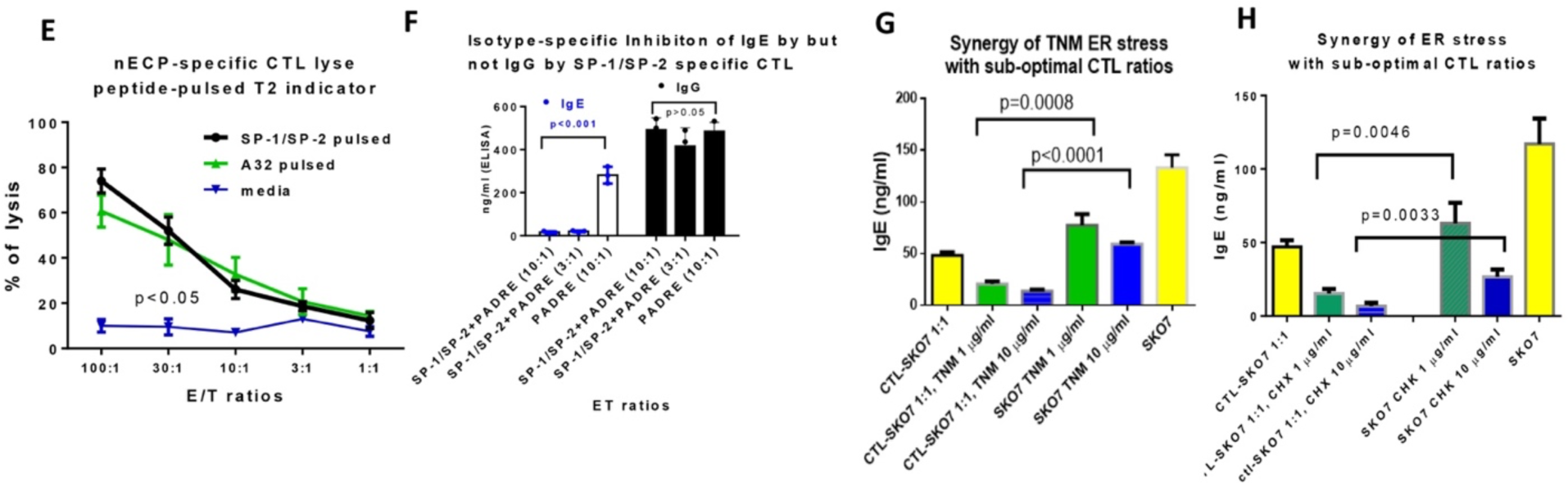
Determination of natural IgE cytotoxic peptides (nECPs) as universal allergy vaccines (UAVs): the correspondence principle. **A.** Upregulation of surface HLA-B7.02: nECP candidates were screened using HLA- B7.02 gene transfected TAP-deficient T2 cells (a gift of Dr. C.T. Lutz) ^14^, not capable of surface HLA-B7.02.01 expression in the absence of its binding peptides. Synthetic peptides according to HLAB7.02 binding motifs of IEBD and MHCPan4.0 were prepared. T2 cells were treated with 10 μg/ml with the following candidate peptides. In the upper panel, the MFI was shifted with SP1 (HPHBPRALM), and SP2 (NPRGVSAYL) nonamers as well as positive control EBV B7 (RPPIFIRRL) in contrast to SP-4 (LPRALMRST) as the negative control, using anti-HLA-B7.02 antibody (BioLegend, Cat: 372402) by FACS analysis using B-D LSR-II. In the lower panel, SP-- 5, -6, -11 nonamers (sequences not shown) did not shift fluorescence intensity comparable to the negative control SP-4 (shown twice) in a second screening. **B.** PBMCs from HLA-A2.01 donors were cultured in 50 ng/ml FLT3 L and stimulated with 0.5 µg/ml ssRNA40, 10 µg/ml A32 peptide (VTLGCLATGY), and 2 µg/ml promiscuous PADRE helper peptide (AKFVAAWTLKAAA) ^15, 16^ and were compared to stimulation using ELA-10 (ELAGIGILTV) with PADRE as positive control. Cells were restimulated with HLA-A2^+^ HLA-A2.01+ EBV B cells, EBV-JY*, pulsed with A32 or ELA-10 without PADRE, treated with brefeldin A for FACS analysis. **C/D.** PBMCs (HLA-A2.01+ and B07.02+) were stimulated with SP-1, SP-2, with PADRE with/without an AMPK mix consisting of AICAR mixture (1 mM) and metformin (50 μg/ml), and then restimulated on Day 10 for analysis by FACS. **E.** Cytotoxicity assay: PBMCs stimulated with A32-, SP-1-, and SP-2 with PADRE, or media treated MoDCs as control ^70^. Cultures were restimulated on day 7 with anti-CD40-treated 32- or SP-1/SP-2 pulsed T2 ^14^ overnight. Re-stimulated CTLs were incubated with 5x10^4^ β2-microglbulin-treated A32- or SP1/SP2-pulsed T2 vs. medium-incubated control in an LDH assay at different E/T ratios (Thermo Fisher kits)^14, 71^. **F.** IgE/IgG inhibition: Day 7 PBMCs primarily stimulated with nECP-PADRE pulsed MoDCs or PADRE-pulsed MoDCs were mixed with 1x10^4^ IgE-producing SKO7 (open bar, typed: HLA-A: 02.01.01G, 03.01.01G; HLA-B: 07.02.01G, 40.01.01G), or IgG-producing EBV-transformed EBV JY* (filled bar, HLA- A2.01, HLA-B7.02, a gift of J, Sidney and A. Sette, LJI) at 10:1 or 3:1 E/T ratios for 24 h, and human IgE and human IgG levels were determined by ELISAs. Four replicate samples were measured in each group. The mean and standard deviation were calculated, and a parametric two-tailed Student’s t test was performed; p values are indicated. **G/H.** SKO7 cells were treated with cycloheximide (CHX) or tunicamycin (TNM) at 1 to 10 μg/ml overnight, washed, and then added to Day 7 primary CTLs as above at the suboptimal 1:1 (E/T) ratio to detect synergy by ER stress primed SKO7 as targets. The day 2 supernatants were collected for IgE ELISA.

Next, it is of critical importance to demonstrate that A32 or SP-1/SP2-MHCI restricted CTL have natural targets in IgE antibody system. Herein, we propose the hypothesis that nECPs correspond to nECPs as a component of MHCIa on IgE B cells are commensurate with the cytotoxic vaccine candidate epitopes as universal IgE allergy vaccines. To induce CTLs, we tested whether disrupting the tolerance of CTLs to nECPs with a helper peptide leads to the induction of nECP-specific CTLs that inhibit IgE production by IgE B cells polyclonally activated regardless of allergen specificities as an UAV candidate. We showed that SP-1, SP-2, and A32 IgE peptides fulfill the criteria of not only exert cytotoxicity using indicator cells artificially pulsed with target peptide but also lyse or inhibit IgE production on IgE producing cells exhibiting the corresponding naturally processed IgE peptides decorated on surface MHCI as natural targets. nECP-specific CTLs were prepared by stimulating with A32, vs. SP-1 and SP2 with PADRE or PDDRE alone, according to the method of Appay ^5, 16^ as described above (Fig. 1B). Fig. 1E showed that SP-1 and SP-2 tetramer-specific and A32-primed and boosted CTLs, but not media-treated control efficiently lysed nECP peptide-pulsed HLA-A2.01 and HLA-B7.02 matched T2 indicator cells. Moreover, to demonstrate the immunogenicity of nECPs exhibiting natural target specificities as an actual vaccine, Fig. 1F showed that SP-1 and SP-2 tetramer-specific CTLs, inhibited ∼95% human IgE production by HLA-A2.01+/HLA-B7.02+ IgE-secreting SKO7 myeloma cells ^10^ at equally at the 3:1 or 10:1 E/T ratios. Moreover, cultures incubated with PADRE, costimulatory signal alone did not inhibit IgE production against SP-1/SP-2 expressing SKO7. Furthermore, the lytic and inhibitory activities are stringently restricted to the human IgE isotype, and nECP-specific CTLs, nor PADRE alone-treated PBMCs did not inhibit human IgG production by human IgG-secreting HLA-A2.01+ HLA-B7.02+ EBV-JY*A B lymphoma cells (provided by J. Sidney and A. Sette, LJI, La Jolla, CA). (Fig. 1F). The lifespan of plasma cells that produce IgE at a high rate is regulated by ER stress ^23^. The inhibition of IgE production by SKO7 at the suboptimal 1:1 E:T ratio was synergized by treatment with the ER stress inducers cycloheximide (CHX) (Fig. 1G) and tunicamycin (TNM) (Fig. 1H) ^23^. These results indicate that the internal ER stress, required for nECP binding to MHCIa supertypes on SKO7 IgE plasma cells, also contributes to the MHCIa supertype-restricted CTLs (synthetic human IgE peptides tested were listed in Suppl. Table 1 to 3).sensitivity of targeted lysis by nECP-

### 2. Abrogating human IgE production by SKO7 and reversal by drug treatment, and dynamics of conversion of A32-stimuated CD8+ T cells to TGF-β and FOXP3 CD4+ T cells

Next, we explored the kinetics and the potency of IgE suppression of IgE myeloma SKO7 using FLT3L treated PBMCs ^15^ stimulated with the respective nECP as a monovalent vaccine: A32 (restricted to HLA-A2.01), or SP-1, or SP-2 (restricted to HLA- B7.02) or a combination of SP-1/SP-2 concomitant *with the helper peptide PADRE, or PDDRE alone as control.* Figure 2A showed CTLs induced by the respective nECP monovalent vaccines, characterized in Fig. 1 potently inhibited human IgE production by SKO7 myeloma cells (HLA-A2.01+/HLA-B7.02+) profoundly to baseline levels at the rapid onset of 8h, lasting 24 h and up to the entire 48 h short-term culture ^11^. Noticeably, each respective monovalent vaccine is equally potent, thus the purpose of combined usage is primarily for encompassing the overall patient population.

**Fig. 2.**
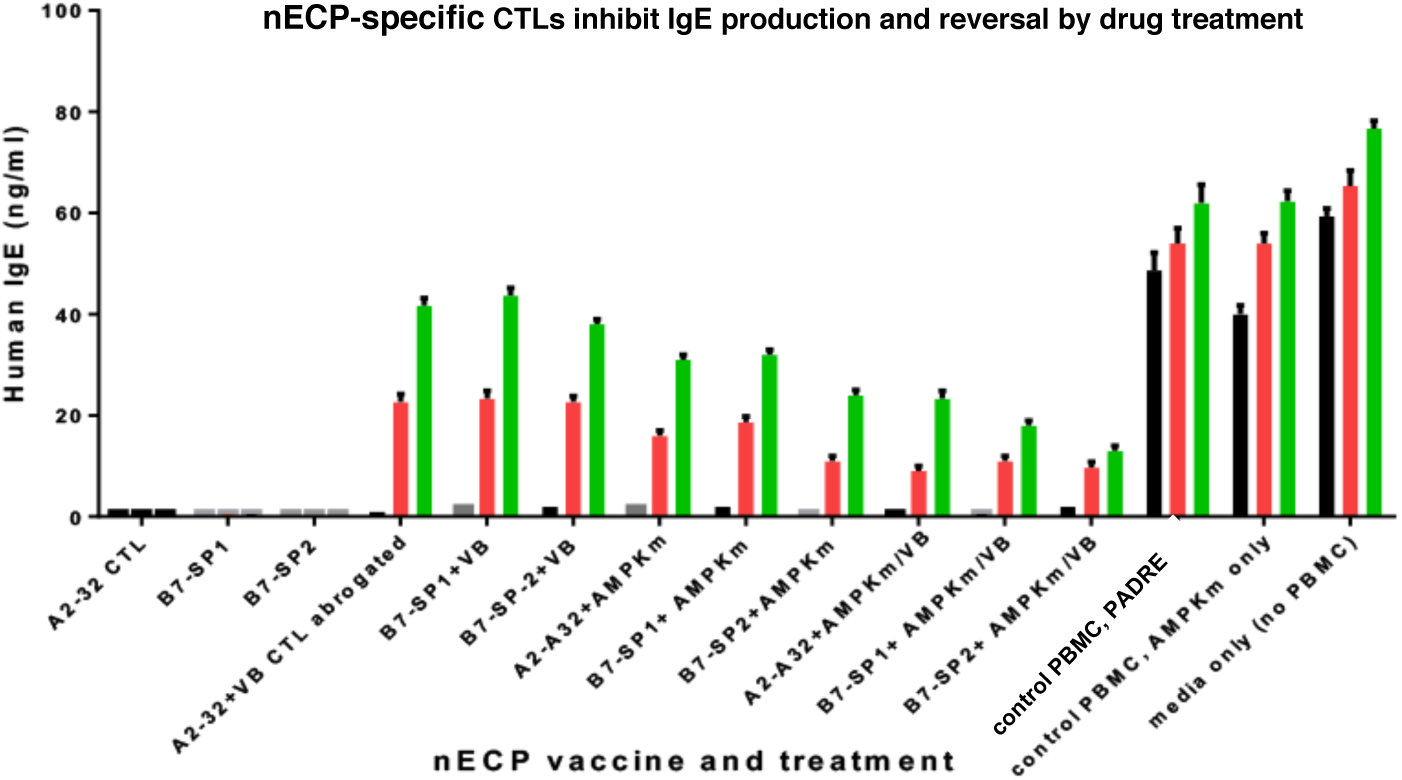
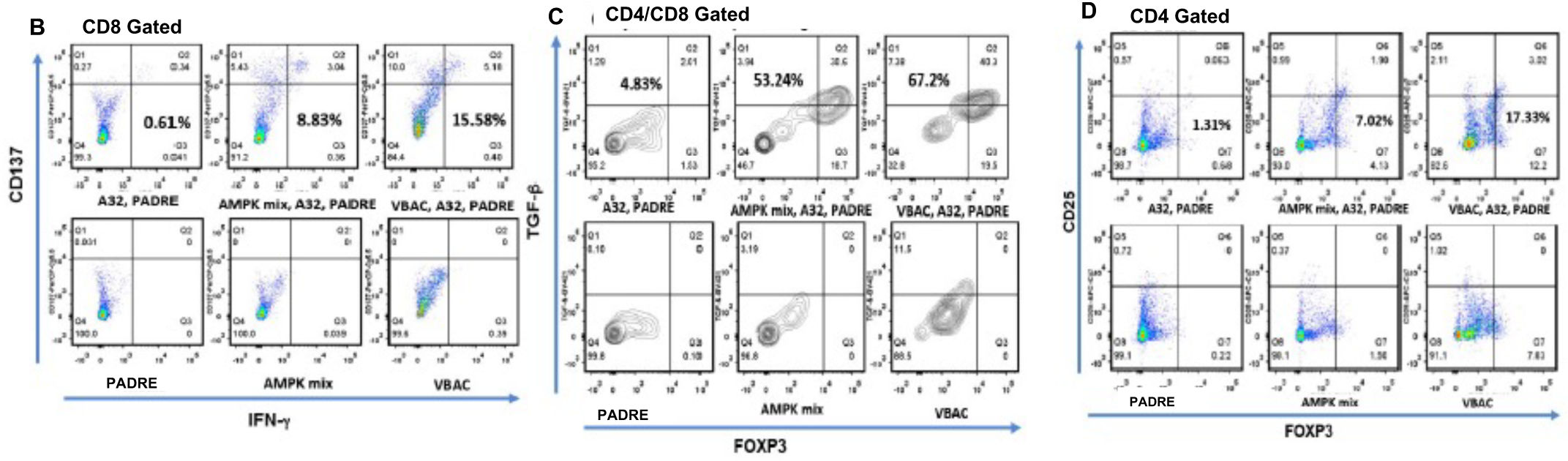
Abrogating human IgE production by SKO7 and reversal by drug treatment and dynamics of conversion of A32-stimuated CD8+ T cells to TGF-β and FOXP3 CD4+ T cells (newly added) **A**: PBMCs of HLA-A2.01 and HLA-B7.02 expressing donors were *in vitro* immunized with nECPs of A32 or SP-1/SP2, PADRE promiscuous helper peptides (PHP) with or without AMPK mix (AICAR + metformin); or VBAC: calcitriol (VD3; 100 nM), bexarotene (Bexa; 100 nM), and vitamin A (retinoic acid) (100 nM); or VB: (VD3; 100 nM), bexarotene (Bexa; 100 nM) and subsequently incubated with SKO7 at a 10:1 ratio, and supernatants assayed by an IgE ELISA. Quadruplicates were set for each group and experiments repeated five times by different individuals. Two-tailed t-test was performed. PBMCs containing CTLs from the above different treatment groups were mixed with SKO7 cells at the optimal ratio of 10 CTL:1 SKO7 in a 200 µl culture in complete RPMI1640. Fifty µl supernatant was harvested at 6 hours (black bar), 24 h (red bar), and 48 h (green bar) successively by spinning the plate at 700 RPM for 7 min at rt followed by multichannel pipette harvest for each period, gently resuspended for further coincubation. Supernatants collected at each time points were prepared for the total IgE ELISA assay. **B**: PBMCs cultured PBMCs cultured using the modified Appay method ^15, 16^, were stimulated with A32 peptide (10 μg/ml) and PADRE (10 μg/ml) in the presence of AMPKm and VBAC calcitriol (VD3; 100 nM), bexarotene (Bexa; 100 nM), or vitamin A (retinoic acid) (100 nM). In contrast, PBMCs incubated in media or in AMPKm and VBAC modulator alone were initiated. On day 10, cells were fixed in 200 µl 1% PFA for membrane and intracellular staining, permeabilized with 200 µl 0.1% Triton X-100 for nuclear staining with fluorochromes-conjugated mAbs by FACS. CD4+ population was analyzed for FOXP3 (AF488), CD25 (APC-Cy7), TGF-ß (BV421); CD8+ population was analyzed for IFN-γ (APC), CD137 (PerCP-Cy5.5), CD4 (BV570) and CD8 (PE-Cy7).

Since IgE plays a key role in parasite defense, surprisingly, AMPK activators mix (AMPKm): AICAR and metformin ^24-27^; or VB: vitamin D3 (VD3) plus bexarotene (an RXR agonist) ^28, 29^ restored IgE production up to 69% apparently by suppressing nECP- UAV-specific CTLs. As shown Fig. 2A, CTL activities against IgE-secreting SKO7 remained intact during the first 8r drug treatment using AMPKm or VB (black bar), and the activities were partially lost, accompanied by reversal of IgE production by SKO7 from 24% to 56% at 24 h (red bar), and 41% to ∼69% at 48 h (green bar) as compared to comparable IgE production among three types of controls: SKO7 incubated with PADRE-treated PBMCs, with media-treated PBMC with drugs, or with SKO7in in media alone. Interestingly, restoring IgE production appeared more effective with treatment of the single AMPKm, or VB, as compared to that of combined AMPKm and VB. In summary, the observations indicate that FLT3L-assisted nECP-stimulated CTLs exhibited two diametrically different functional states, capable of inhibiting IgE production by SK07, and that capable of restoring IgE production by SKO7 following drug treatment.

Notably, nECP A32 vaccine was discovered not by upregulation of surface HLA-A2.10 in TAP1/2 deficient T2 cells (Fig. 1B). Indeed, there were multiple unproductive attempts in preparing A32-HLA-A2.01-PE tetramers through efforts of NIH tetramer facility and contracted commercial service. This is likely due to the low affinity binding of A32 also in upregulate surface expression in the TAP1/2 deficient T2 cells. Nevertheless, A32 is demonstrated a potent vaccine candidate via direct DC-based immunization. Because of its importance, FACS analysis was conducted for the frequencies and dynamics using cytokine biomarkers for the A32-HLA-A2.01 specific T cells, activated by A32 presenting FLT3L derived cDCs, with/o drug treatment in Day 7 PBMC cultures.

Fig. 2B showed CD137+ (4-1BB)/IFN-γ+ double positive CD8+ T cells were elevated to 0.61% with PADRE costimulation, which correlated with its potency to suppress IgE production by SKO7 cells (Fig. 2B). In contrast, double cytokine positive CD8 T cells were undetectable in cultures stimulated with PADRE costimulation alone (∼0% in media). Despite the increase of CD137+/ IFN-γ/ T cells (8.83-15.5%, shown in Fig. 2B due to AMPKm or VB treatment, these cells failed to inhibit IgE production by SKO7 cells, or paradoxically restored IgE-production by SKO7 (Fig. 2A).

Noticeably, Fig. 2C showed that ∼53.2 to 67.2% of the transitional double positive DP CD4+CD8+cells, expressing TGF-β and FOXP3 were present in the AMPKm or VB- treated, A32-, PADRE-stimulated cultures. Fig. 2D showed high frequencies (7.02- 17.33%) of A32-activated FOXP3/CD25 CD4 T cells in 7.02% AMPKm-treated, A32- stimulated cultures as well as 17.33% in VB-treated, A32-stimulated cultures. The increment of A32-specific T cells with PADRE in the metabolic milieu was compared to low to non-detectable levels of the respective cell type in cultures treated with PADRE costimulation alone, or those treated with drugs alone (either type). The observations suggest that the transformed FOXP3/CD25+ A32-specific of Treg by FLT-3 cDCs may counteract A32-activated CTLs, which will be investigated using normal PBMCs as the physiological target besides the potency observed using IgE myeloma cells,

### 3. Triple transgenic chimeric human IgE, HLA-A2.01, HLA-B7.02 mice to test the efficacies of nECP universal allergy vaccine

Next, we examined whether nECP vaccines inhibit human IgE production *in vivo*. The correspondence principle of nECP vaccines, are shown in triple transgenic animals, capable of human IgE production, expressing natural nECP decorated onto human HLA-A2.01 and HLA-B7.02 via the endogenous ER stress, as well as inherent of self-tolerance to human IgE. The breaking of human IgE tolerance and inhibition of human IgE can then be tested by vaccinating nECP vaccines in the rodent model. Fig. 3A and Fig. 3B showed that triple transgenic mice backcrossed to C57B/6 nine generations expressing chimeric NP-specific human IgE ^30^, HLA-A2.01 and HLA-B7.02 (IGE-A32-B7-114) were established and maintained in the animal facility. Nitrophenol rodent (NP)- specific VH is fused to human IGE heavy chain (CHε2-4) ^30^ as cDNA constructed under the promoter, EF-1a, wherein the expression of the chimeric antigen-specific chimeric human IgE (base pair: bp 1732-3432) permits the processing of endogenous A32-SP-1-SP2 nECPs via ER stress in transgenic B cells. Moreover, human HLA-A2.01 (bp 5795- 8907) and HLA-B7.02 (bp 4509-5611) cDNA fused through P2A cleavable sites were constructed under the CMV promoter in the reverse orientation, therefore the processed nECPs were decorated on HLA-A2.01 and HLA-B7.02 of transgenic B cells as targets using the triple transgenic vector (DNA sequences of the three transgenes were listed in Suppl. Fig.1). Noticeably, the distinct advantage of the triple transgenic mice is the establishment of central nECP tolerance to HLA-A2.01 and HLA-B7.02 during immune ontogeny, and the breaking tolerance by providing the second promiscuous measles helper peptide non-cognately via liposomes as a surrogate animal model for human vaccines. Founder mice exhibiting triple human HLA-A2.01, HLA-B7.02, and chimeric NP-specific human IgE were selected herein: 2 female founder #30 and #39 and two male founder #26 and #36 with gene identifier PCR fragments for IgE (GHuIGE, fragment of 330 bp); and HLA-A2.01 (GHLA-2.01, fragment of 321 bp); and HLA-B7.02 (GHLA-B7.02, fragment of 324) against the molecular weight marker (M) and internal molecular tag (MT) amplified from the IGE-A32-B7-114 dual promoter vector construct (Fig. 3 C-E) identified for the four founder mice for establishing the mouse colony.

**Fig. 3.**
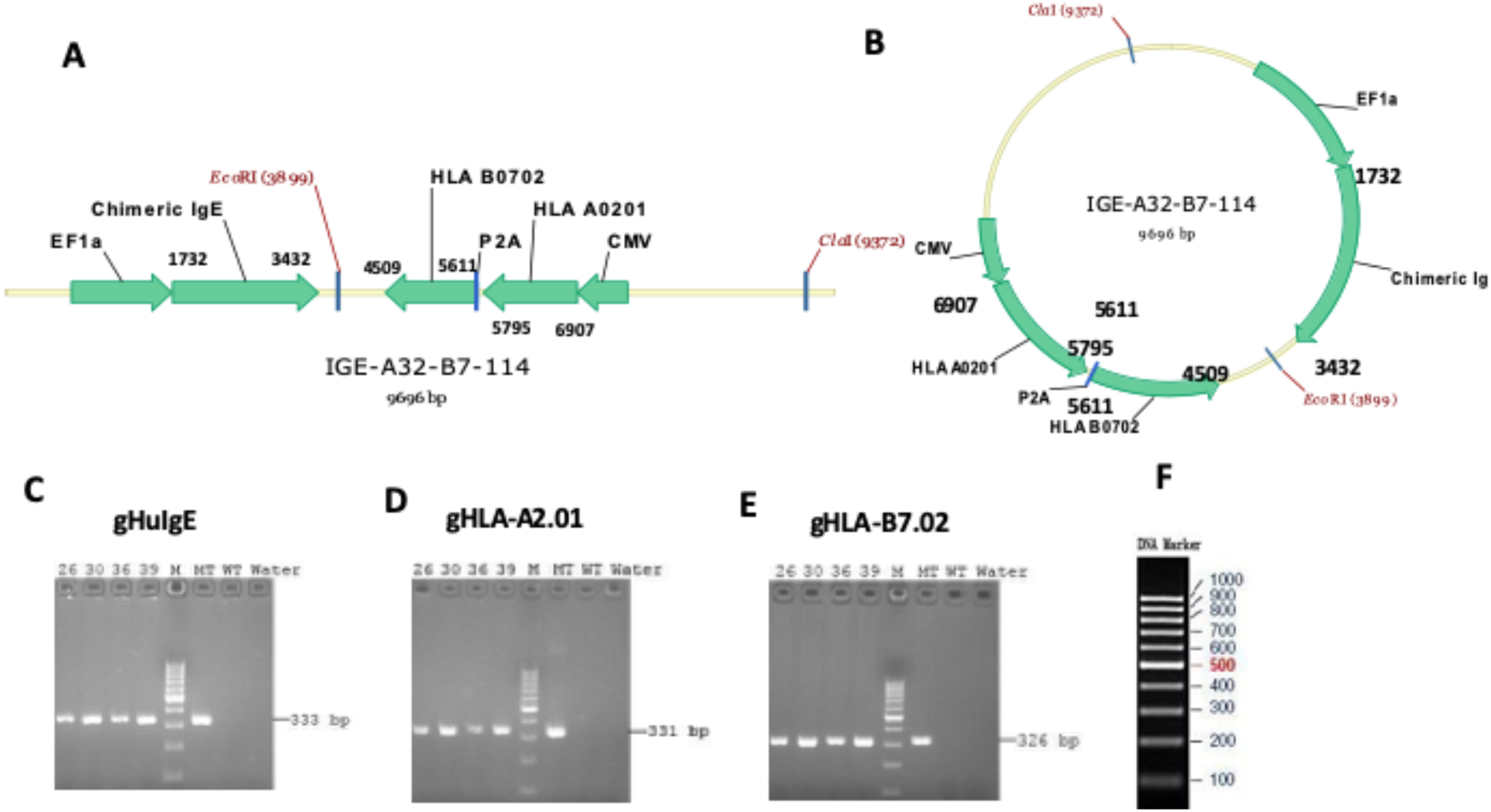
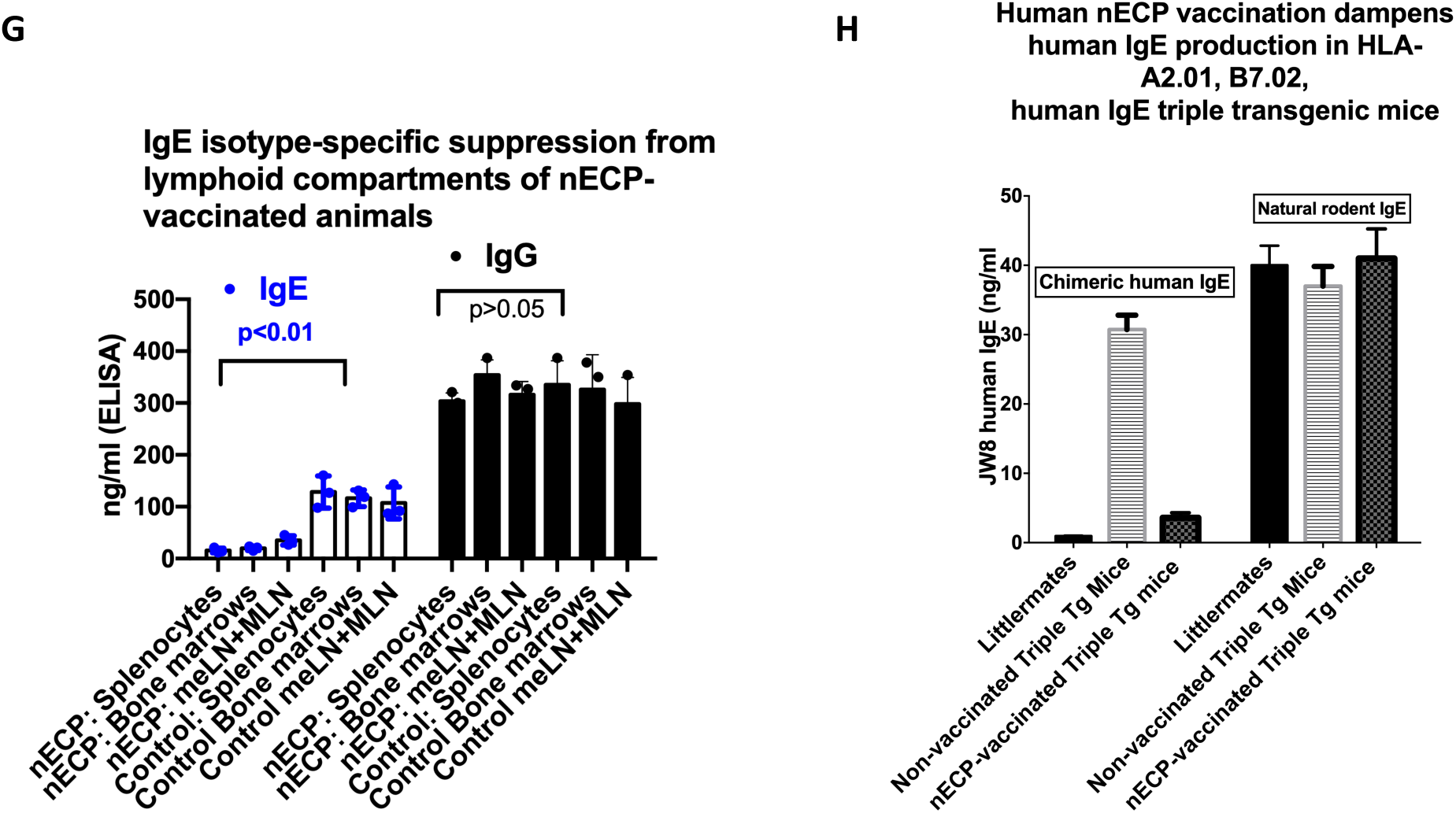
nECP-specific HLA-A2.01- and HLA-B7.02-restricted CTLs dampen natural chimeric IgE levels *in vivo*. The triple Tg mice, backcrossed for nine generations were housed in an OLAW-licensed ARF at the Institutions. **A/B.** Triple transgene construct of HLA-A2.01/HLA-B7.02 and chimeric JW8 human epsilon heavy chain under the respective CMV and EF1a promoter control. **C-E.** Diagnostic PCR fragments of HLA-A2.01, HLA-B7.02, and the JW8 heavy chain of the founder mice. Founder mice exhibiting triple human HLA-A2.01, HLA-B7.02, and chimeric NP-specific human IgE were selected herein: 2 female founder #30 and #39 and two male founder #26 and #36 with gene identifier PCR fragments for IgE (GHuIGE, fragment of 330 bp); and HLA-A2.01 (GHLA-2.01, fragment of 321 bp); and HLA-B7.02 (GHLA-B7.02, fragment of 324) against the molecular weight marker (M) and internal molecular tag (MT) amplified from the IGE-A32-B7-114 dual promoter vector construct (Fig. 3 C-E) identified for the four founder mice for establishing the mouse colony. Genotyping of the Tg mice. Genomic DNA was prepared from mouse tails using a genomic DNA preparation kit. IgE, HLA A0201, and HLA B0702 were measured by PCR with the primers CTTCCATTTCAGGTGTCGTG (forward) and CAAGCTGA TGGTGGCA TAGT (reverse) for IgE, CACGTAGCCCA CTGCGATG (forward) and CAATGGGAGTTTGTTTTGGCACC (reverse) for HLA A0201, and CTACGACGGCAAGGATTACAT (forward) and GGGCAGCTGTGGTGGTGC (reverse) for HLA-B07.02. The PCR conditions were 35 cycles of 94 °C for 20 s, 60 °C for 30 s, and 72 °C for 35 s. **F.** Tg mice and littermates (group of 5 mice each) were vaccinated with 20 μg each of A32, SP-1, and SP-2 with 10 μg MHP (ISISEIKGVIVHKIETILF) in 250 μl, mixed in 250 μl DOTAP, and primed/boosted via subcutaneous injection at 10-day intervals according to the previously published protocol ^10^. Sera collected on Day 7 after another reboot were evaluated by human and rodent IgE ELISAs. ELISAs for measuring human IgE were performed using an HRP-conjugated anti-human IgE antibody (Bethyl Inc., TX). Rodent IgE was measured using DNP-specific hybridoma IgE as a standard, and rat mAb anti-IgE was made by this laboratory ^4, 72, 73^. **G.** Spleens, bone marrows ^74^, and mucosal mesentery and mediastinal lymph nodes (MLN/MeLN, pooled from 5 mice) were prepared on Day 7 after the booster and mixed at 10: 1 E/T ratios with human IgE producing HLA-A2.01+/HLA-B7.02+ SKO7 as well as human IgG producing HLA- A2.01+/B7.02+EBV JY*A cells (gift of J. Sidney and A Sette at LJI, La Jolla, CA). The day 2 supernatants were measured by ELISA for human IgE and IgG. Mean and standard deviations were calculated, and a two-tailed Student’s t test was performed with the respective treated mice vs. control mice. CTL assays were performed as described (Fig. 1E legend) ^72^.

Mice were primed and boosted with triple A32, SP-1, SP2 nECPs, with measles promiscuous helper peptide (MHP) ^31^ for the consideration of herd immunity. The nECP vaccine was administered in DOTAP liposomes subcutaneously, presented DCs, and the extent of CTL activation was commensurate with bone marrow-derived dendritic cells (BMDCs)-based immunization as described in the laboratory ^10^. As shown in Fig. 3F, triple Tg mice produced a natural level of NP-specific chimeric human IgE, which was dampened in nECP-vaccinated mice but not in littermates. In contrast, rodent IgE levels were not affected, showing that the effect of the human nECP vaccines strictly inhibited chimeric human IgE production without cross-inhibiting that of endogenous rodent IgE produced normally by the triple Tg animals; serum IgG levels in both Tg and normal mice or littermate (not shown) indicating nECP-vaccination downregulates human IgE as a class uniquely without affecting rodent IgE or IgG.

Next, we wish to ascertain whether nECP-specific HLA-A2.01 and HLA-B7.02 CTLs from the triple Tg mice can likewise inhibit human IgE production by human IgE producing HLA-A2.01/HLA-B7.02+ SKO7 myeloma cells due to match of nECPs as well as MHCI. As shown in Fig. 3G, single cells prepared from the spleens, lymph nodes, and bone marrow of nECP-vaccinated triple Tg mice inhibited approximately 75-92% of human IgE production by SKO7 myeloma cells but did not affect human IgG production by EBV-JY*A transformed B cells for a different human isotype. Thus, downregulation of transgenic human chimeric JW8 IgE-producing B cells and plasma cells *in vivo* in nECP-vaccinated triple Tg mice was strictly concordant with that in human IgE-producing myeloma cells but not a different immunoglobulin isotype. Since all human IgE B cells and plasma cells express a single isotype of IgE heavy chain regardless of allergen specificities, the inhibition of chimeric IgE production in triple transgenic Tg rodent model strongly suggest that the nECP vaccines may be used to treat diverse, allergen-specific, e.g., pollen, cockroach, house dust mite, food, or peanut allergens, IgE-mediated allergic inflammation.

### 4. Cellular Mechanism: The natural target of nECP vaccines and homeostasis of nECP-specific Tregs: the three signals for Self/Non-Self/Self

Next, it is critical to examine whether nECP-tetramer-specific CTLs can directly inhibit and correct overproduction of IgE in normal human PBMCs *ex vivo*. it is pertinent to investigate mechanisms whereby IgE responses may be fully recovered in the chemical milieu as described in Fig. 2A-B; therefore upon removal of IgE danger signals, IgE-mediated immune defense may be restored. For this purpose, the SP-1/SP-2 tetramers-specific T clonotypes can be conveniently monitored in lymphocyte subsets of human PBMCs.

Thus, a three-stage human PBMC culture systems was developed to test inhibition of IgE production by human PBMCs and its homeostatic recovery. PBMCs of more than 50 separate donors are collected from the local San Diego Blood Bank (SDBB). Donation was not repeated. Except demographic data, the identity and IgE allergic status are protected (Source of PBMC detailed in Suppl.). Single or double HLA-A2.01 and/or HLA-B7.02 after genotyping were employed in the studies. First, FastDCs (or MoDCs) were pulsed with SP-1/SP-2 peptides/PADRE. In stage 2 cocultures, nonadherent T cells were added to nECP-pulsed DC cultures, wherein different immune response modifiers were added at different days of incubation for inducing tetramer-specific CD8+ CTLs or converting to tetramer-specific CD4+ Tregs. Next, in stage 3 cocultures, fresh human IgE producing PBMCs, stimulated with anti-CD40 and IL-4 from the same donor were added to activated nECP-specific CTLs of stage 2 cultures for dampening and recovery of IgE responses (stepwise methods and execution scheme was detailed in Method of Suppl.). Notably, the four salient features are: First, Figure 4A showed that SP-1/SP-2 stimulated CTLs profoundly inhibited (> 90%) IgE production in stage 3 cocultures in normal media as positive controls. This observation crucially establishes the immune-physiological role of nECP-specific CTLs in inhibiting the hyper-IgE production extended to normal human PBMCs according to the principle of correspondence. Figure 4B showed that neither nECP-specific CTLs nor nECP-specific CD4 T cells negatively affected IgG production, indicating the exquisite specific IgE isotype-specificity without cross reactivities. IgE recovery plays a key role in the IgE-mediated parasitic defense that ensures the safety of IgE-mediated immune defense of the nECP-UAV vaccine ^32^. We propose that immune homeostasis is influenced in the context of energetics and metabolomics by the endogenous AMPK energetics and nucleic receptors (NR). We therefore investigated whether nECP-specific CTLs may be abrogated or converted into immunoregulatory T cells (Treg) treated with AMPK activators ^21, 22, 25, 33^, or NR agonists for vitamin D receptor, (VDR, calcitriol) ^28^, retinoid X receptor (RXR, bexarotene) ^29^ e.g., VB, or PPARα (oleoylethanolamide, OEA) ^34^. As shown in Fig. 4A, the capacity of CTLs to inhibit IgE production by PBMCs were abrogated by nonadherent T cells stimulated with nECP pulsed AMPKm treated MoDCs from day one during induction up to day 7 day as effectors in the stage 2 DCs/T cell cocultures. In contrast, VB was effective at culture initiation but not added at later time. Restoration of IgE production in the nECP-stimulated, drug-treated stage 3 cocultures was comparable to control stage 3 cocultures of PBMCs added with media-treated DCs/NAD or PBMC in media alone.

**Fig. 4.**
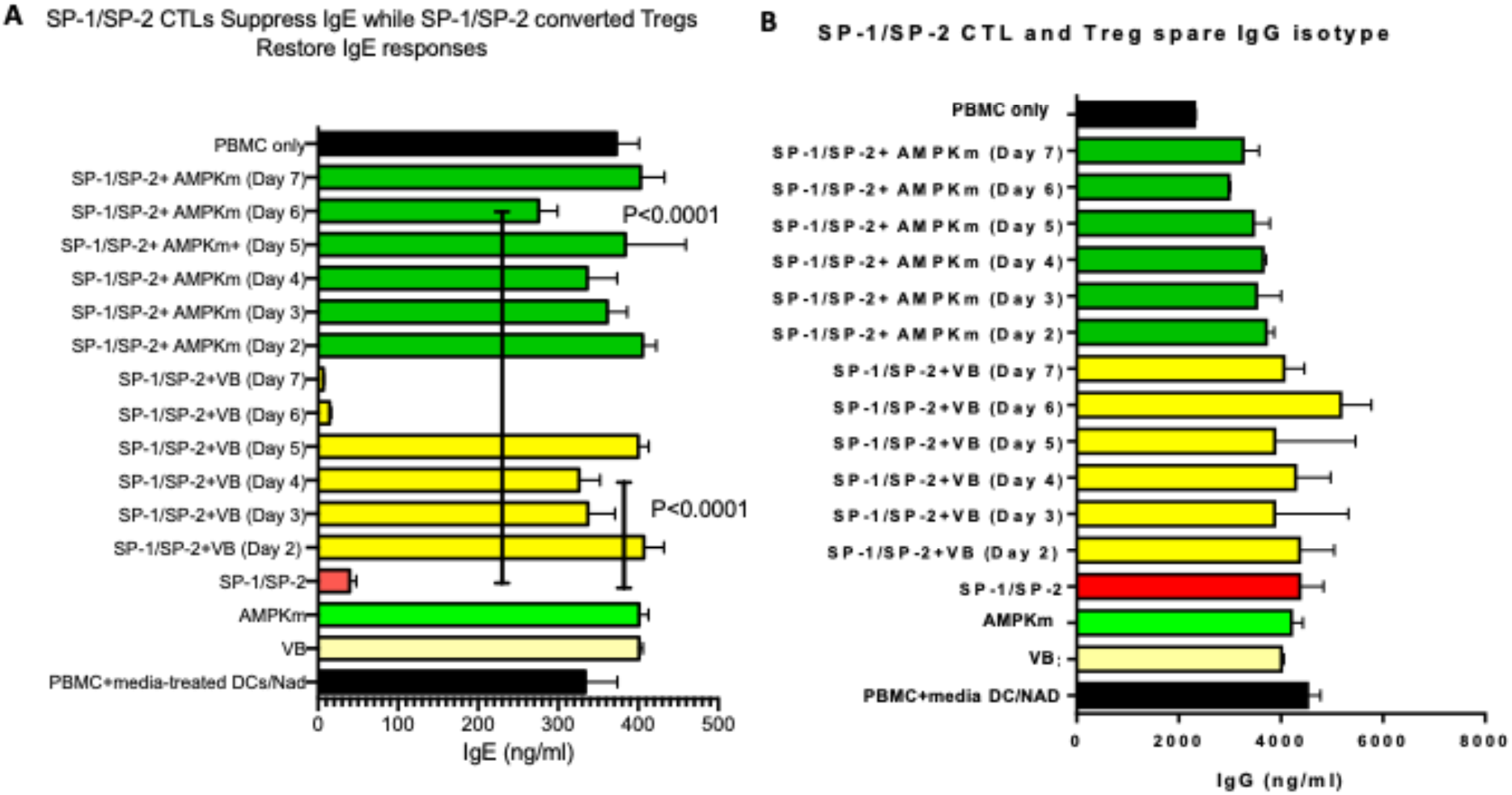
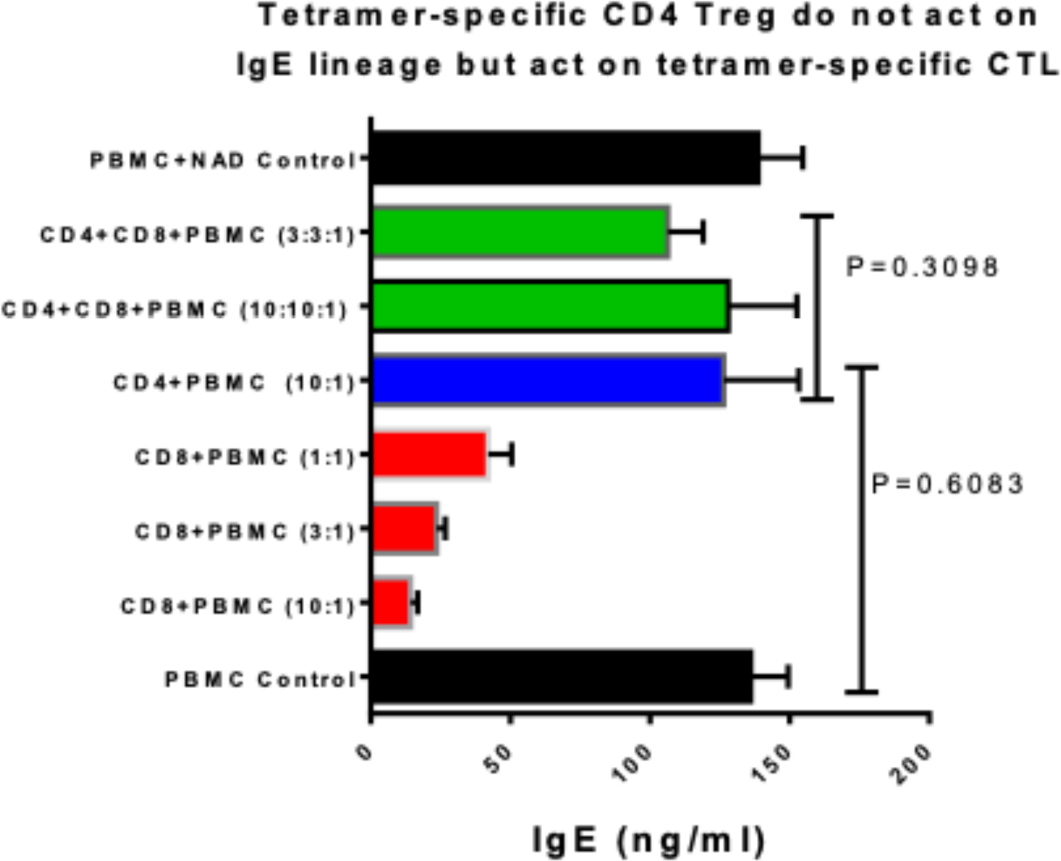
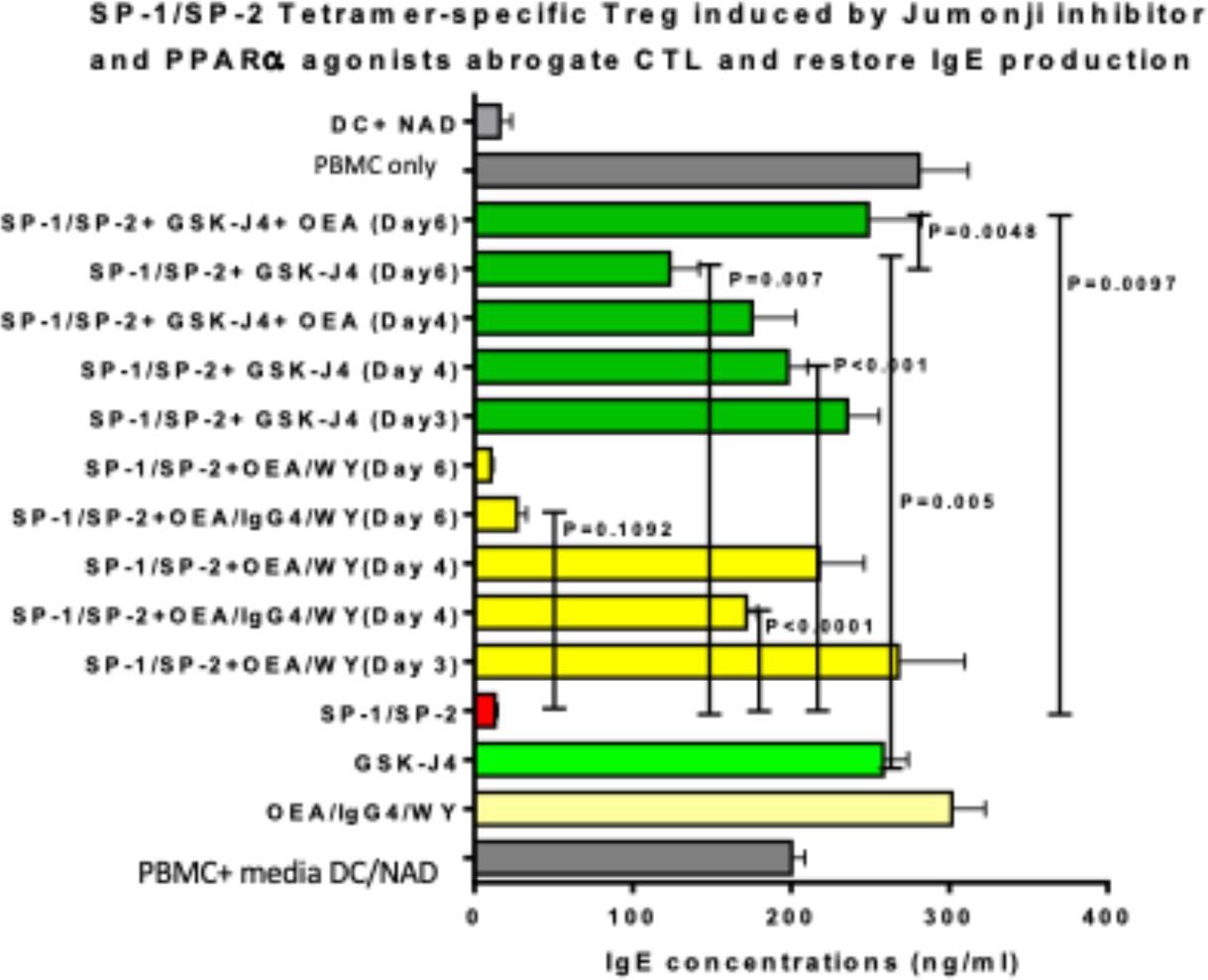
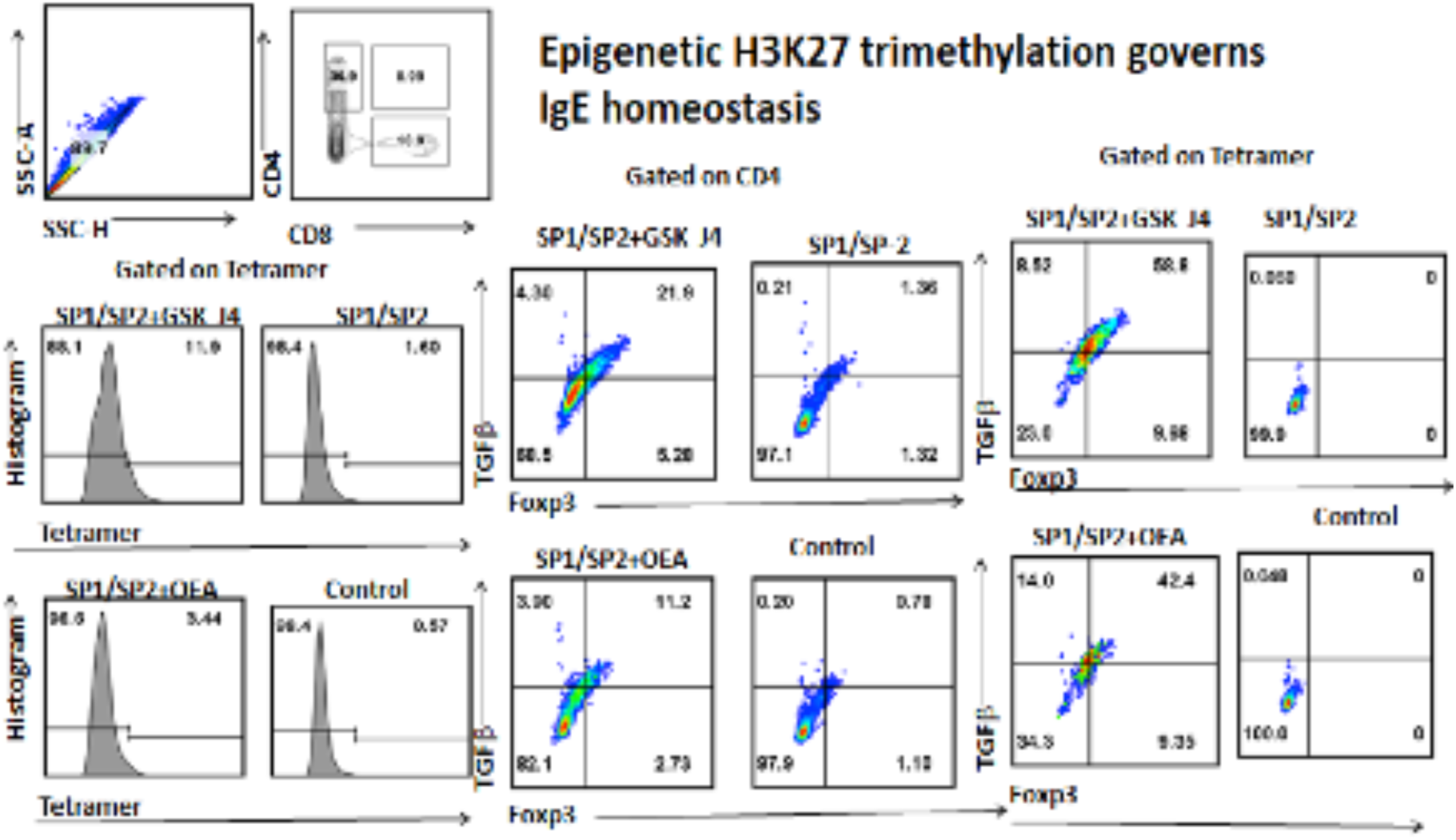
Regulation of the IgE producing cells: the homeostatic principle. PBMCs from healthy donors from the San Diego Blood Bank (SDBB), are first stained for surface HLA-A2 (mAb BB7.2) and HLA-B7 (mAb BB7.1, BioLegend, Inc.), and the HLA-A2.01 and HLA-B7.02 were evaluated by PCR genotyping (Histogenetics, N.Y.). Individuals double or single positive for HLA-A2.01, and/or HLA-B7.02. were used throughout the coculture experiments. The demographics and details with contract SDBB are described in Supplements. **A/B.** FastDCs ^75^ were prepared from PBMCs convenient for tetramer analysis, and pulsed with SP-1/SP-2 and PADRE. Stage 2 cocultures were initiated by adding NAD 3 X10^5^ (FastDCs: NAD = 1:3) in quadruplicate with/o the AMPK mix (AMPKm) or vitamin D3 or calcitriol (100 nM)/bexarotene (100 nM) (VB) ^17^. In parallel, IgE cultures were initiated in PBMCs of the same donor with an anti-CD40 antibody, IL-4, IL-6, and IL-21. Stage 3 cocultures were then initiated by adding IgE cultures on Day 1 to Day 6 successively to the above stage 2 cultures. The day 15 and day 18 supernatants (counting from the initiation date of the stage 2 cocultures) were assayed for IgE **(A)** and IgG **(B)** by ELISA. Reversal of IgE produced in the drug-treated stage 3 cocultures was compared to stage 3 coculture of PBMC with media-treated DCs/NAD control as well as PBMC alone control. **C.** The protocol for the three-stage cultures followed the description above except that the stage 1 FastDCs were treated with OEA (30 μM), WY (30 μM) or GSK-J4 (30 μM), and the effects on the IgE lineage were evaluated. Multiple experiments were performed on different types of blood donors, and a two-tailed Student’s t test was conducted with the respective comparison groups. **C.** CD4+ or CD8+ T cells from day 4 stage 2 cocultures were isolated by negative selection. Enriched CD4+ or CD8+ T cells were added to the activated IgE cell lineage, and Day 10 supernatants were assayed for IgE levels by ELISA. **D.** The three-cell model, wherein CD4+ Tregs directly target CD8+ CTLs. **E.** The three cell co-cultures with nECP SP1/SP2 and PADRE with GSK-J4 (30 μM) are described in Panel A. **F.** SP-1/SP-2 tetramer-positive cells were analyzed with fluorochrome-labeled mAb by FACS. More details of the procedures were described in the Supplements.

Next, Figure 4C showed that fractionated nECP-specific CD8 T cells from nECP vaccinated stage 2 cocultures directly inhibited IgE production by anti-CD40 and cytokine-stimulated PBMCs, while CD4 T cells fractionated from nECP-stimulated, AMPKm-treated DCs/T cells cocultures abrogated the capacity of nECP-specific CTLs to inhibit IgE production PBMCs, thereby restoring the IgE production by anti-CD40 and cytokines-stimulated PBMCs. Notably, nECP-specific CD4 Tregs did not suppress IgE producing B cells and plasma cells despite nECPs decorated on HLA-B7.02, indicating a division of labor of nECP-specific Tregs for CTL.

Next, the homeostatic regulation is extended further to an epigenetic model. Epigenetic control is well documented for immune cell lineage development ^35^. The Jumonji (JMJ) family of histone demethylases are Fe^2+^- and α-ketoglutarate-dependent oxygenases, including H3K27me3-specific demethylase subfamily KDM6 subfamily member, JMJD3 that regulates transcriptional chromatin complexes. GSK-J4, a H3K27 JMJD3 demethylase inhibitor plays an anti-inflammatory role in downregulating TNF-α production by macrophages ^36^; therefore we examine its role in generating immunoregulatory DCs. JMJD3 coevolves with the fat mass obesity gene (FTO), which potently regulates glucose vs. lipid metabolism also in the context of metabolic immunology ^37-39^. Figure 4D showed that GSK-J4 treatment in both the early inductive (Day 2) and even as late (Day 6) effector phases in the stage 2 DCs/T cells coculture completely abrogated the capacity of CTLs to inhibit IgE production by anti-CD40 and cytokines-stimulated PBMCs, thus fully restored IgE production in the reconstituted stage 3 cocultures. In contrast, OEA applied alone or when used with WY-14643, a PPARα agonist, inhibited the early inductive but not the late effector phase of CTLs (Fig. 4D).

GSK-J4, the JMJD3 inhibitor-treated stage 1 MoDCs cultures, was tested for transforming tetramer-specific nECP-specific CTLs into CD4 Tregs ^36, 40^. As shown in Fig. 4E, the potency of restoring IgE production was correlated with levels of nECP tetramer-specific FOXP3/TGF-β double-positive CD4 T cells, which were higher in GSK- J4-treated cells (21%) than those in OEA-treated cells (11%) respectively during the transforming SP1/SP2-specific CTLs to SP1/SP2 specific Treg in the stage 2 DCs/T cells cocultures. The observations were reminiscent in that GSK-J4 is as potent as AMPKm in the immunoregulatory generative DCs for transforming the CTLs into Tregs in stage 2 cocultures.

Moreover, the observations coincides with that of Fig. 2D showing that A32 nECP- activated CD4+ FOXP3/TGF-β Tregs were transformed by A32 pulsed, AMPKm-treated FLT-3 stimulated cDCs.

In summary, nECP-presenting generative MoDCs were influenced by microenvironmental milieu, from a diverse nuclear receptors agonists, AMPK activators and epigenetic H3K27 demethylase inhibitors in the stage 1 MoDCs. Then nECP-specific CTLs can be transformed into CD4+ nECP-specific Treg mediated by the generative DCs in stage 2 MoDCs/T cell cocultures. Thus IgE production by PBMCs in the stage 3 cocultures inhibited by enabler DCs/CTLs, are restored by generative DCs/Tregs under metabolic milieu.

### 5. Molecular mechanism of the generative third signal

Next, it is pertinent to explore the molecular mechanisms of the immunoregulatory generative DCs as the foundation of IgE homeostasis. Immunometabolism is linked to effector and immune-regulatory mechanisms of APCs, T- and B-cells ^35, 41^. Recently, Chi and colleagues showed that Treg maintenance is SREBP-dependent ^42^. Herein we showed that at levels of DCs, the three different kinds of drugs for AMPK, nuclear receptors and H3K27 methylation is sufficient for enabling transformation of nECP- specific CTL to Treg. We propose that the above treatments converge in a unifying lipo-oxidation of countering SREBP-1c at the immunoregulatory DCs. Since AMPK downregulates the master lipogenic switch, SREBP-1c transcription factor in favor of lipid oxidation via TSC2 or direct phosphorylated inactivation ^43, 44^, we therefore examine whether AMPK activators, and other drugs employed herein affecting lipid metabolism may converge into inhibitory pathway for SREBP-1c and its downstream lipogenic genes at the levels of the immuno-regulatory MoDCs.

Thus, Fig. 5A-C shows that metformin or metformin/AICAR reduced SREBP-1c protein in MoDCs by more than 90%. GSK-J4, an H3K27 JMJD3 demethylase inhibitor ^36 45^ that potently influence generative DCs shown in Fig. 4F likewise potently inhibited SREBP- 1c expression by approximately 93% in MoDCs (Fig. 5A). As a positive control, CDCA/6E-CDCA (CDCA/obeticholic acid), potent FXR nuclear receptor agonists reduced SREBP-1c protein levels by at approximately 87%. VD3 (including 9-cis) inhibited SREBP-1c protein expression by approximately 85%, which was comparable to the effect of AMPKm (∼79%) (Fig. 5B). Moreover, OEA, which is known to enhance fatty acid β-oxidation ^33^, caused an 82% reduction in SREBP-1c protein levels after 30 min (Fig. 5C). As expected, FASN expression was decreased 2-3-fold after treatment with AICAR, VD3, and OEA as well as conspicuously GSK-J4; moreover, the treatment notably led to the 60 kDa breakdown product of FASN (Fig. 5D). Small heterodimeric partner (SHP), an orphan nuclear receptor (NROB2), plays a critical and ubiquitous role in inhibiting SREBP-1c expression via the FXR pathway ^46^. Similarly, lipogenic genes, including FASN, were downregulated by AICAR and CDCA/6E-CDCA in the HepG2 cell line (Suppl. Fig. 2). We observed AICAR, metformin, and OEA caused a 2.2-3-fold increase in SHP levels (Fig. 5E). Notably, the altered energetics affects the expression of co-stimulatory molecules of MoDCs. CTLA-4 is the key molecule in the CD28 subfamily that maintains self-tolerance against lethal autoimmunity ^47, 48^, and the expression of CTLA-4 was augmented ∼2-fold (1.89∼1.96-fold) in AMPKm- and GSK- J4-treated MoDCs (Fig. 5F). Importantly endogenous FOXP3 upregulated by superantigen B in rodent BMDCs as well as transgenic FOXP3+ human MoDCs were shown to significantly induce Tregs suppressing allergic reaction in vivo and against third party alloantigen in vitro respectively ^49 50^, Interestingly, siRNA knockdown of SREBP-1 concomitantly upregulated FOXP3 expression in MoDCs (Fig. 5G) ^49 50^.

**Fig. 5.**
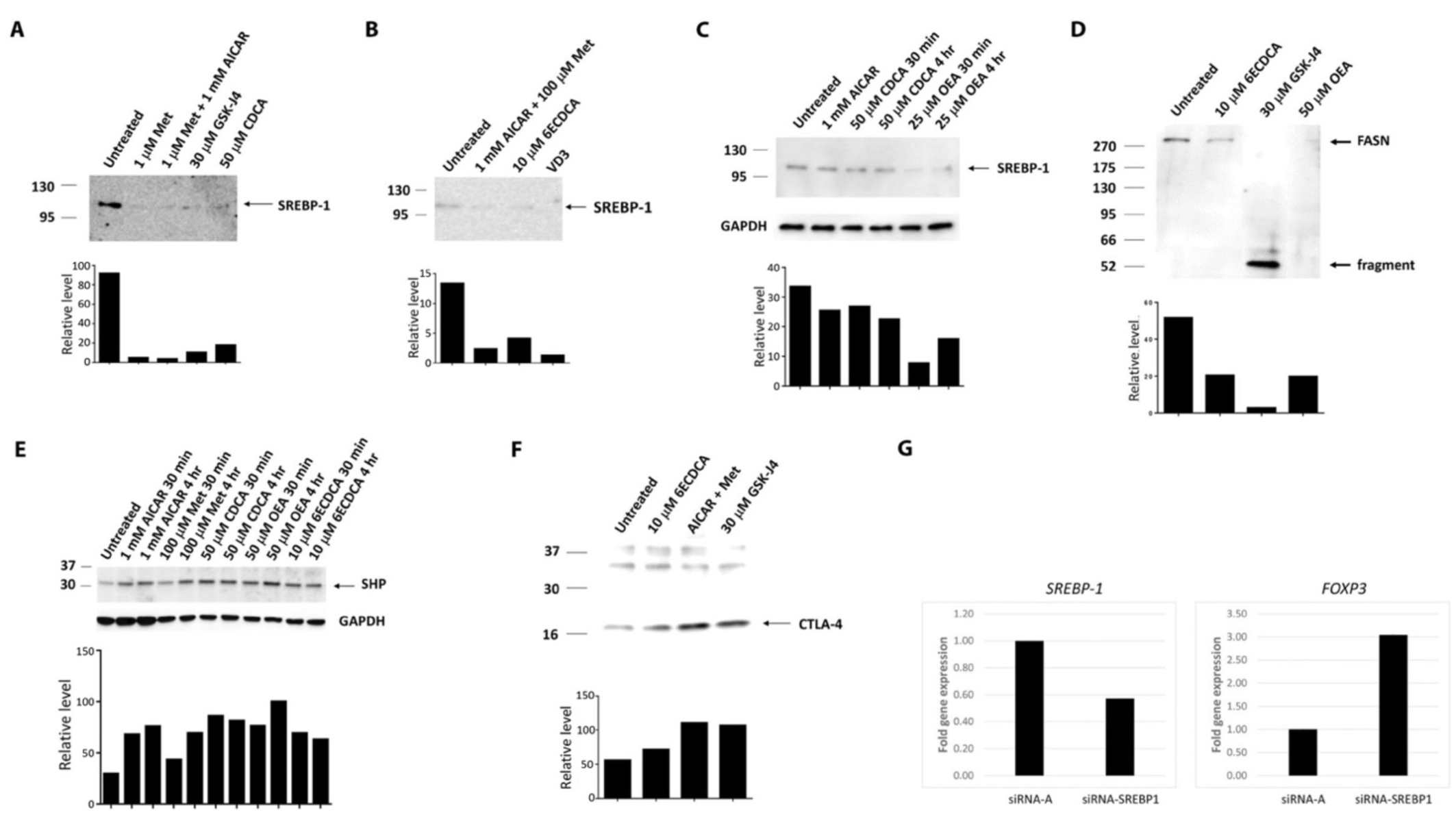
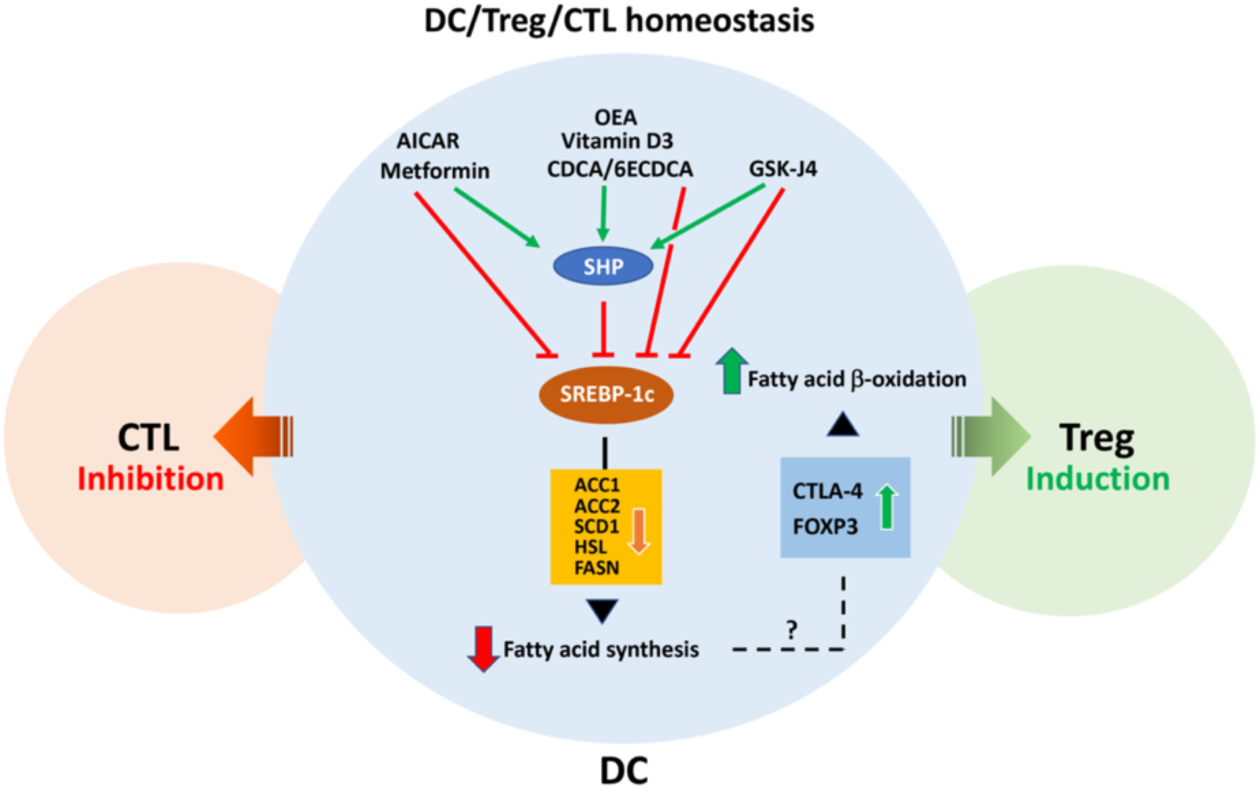
Metabolic milieu of immunoregulation generative DCs. MoDCs were prepared according to the procedures ^70, 75^ described in the Suppl. For drug treatment (A-F), MoDCs were incubated with different drugs at the concentrations indicated. The duration of treatment was 4 hours unless otherwise indicated. Whole-cell lysates were prepared after treatment and resolved by SDS‒PAGE followed by immunoblotting using corresponding antibodies (upper panel), and the relative signal was quantified by ImageJ (lower panel). **A.** AICAR/metformin (Met), CDCA and GSK-J4 downregulate SREBP-1 in immature MoDCs, as shown by immunoblotting (upper panel) and ImageJ (lower panel). **B.** AICAR/Met, 6ECDCA and VD3 (100 nM calcitriol) downregulate SREBP-1 in mature MoDCs by immunoblotting and ImageJ. **C.** OEA downregulates SREBP-1 with others as positive control. **D**. 6ECDCA, GSK-J4 and OEA downregulate FASN in mature MoDCs with 6ECDCA as control. **E**. AICAR/Met, CDCA, 6ECDCA and OEA upregulate SHP in mature MoDCs. **F**. AICAR/Met and GSK-J4 upregulate CTLA-4 in mature MoDCs. **G**. Mature MoDCs were transfected with SREBP- 1c siRNA or control scRNA at a final concentration of 33 nM 24 hours later, and RNA was isolated followed by qRT‒PCR detailed in Supplement. **H.** Homeostatic lipogenic model for generative DCs, explained in text for three signal model.

Taken together, Figure 5H proposes an anti-lipogenic model, based on the observations (Fig. 5A-G) in which preferred lipid oxidation utilized by the immunoregulatory generative DCs in the stage 1 DCs cultures license nECP-specific Treg, transformed from the nECP-specific CD8 CTLs in the stage 2 DCs/T cell cocultures.

## DISCUSSION

### 4.1. Synopsis

We showed that nECPs-specific CTLs were elicited by nECP-UAVs to decrease IgE production targeting the corresponding nECP decorated on MHCIa human IgE B cell lineages encompassing a diverse allergen repertoire. In addition, nECP-tetramer-specific CTLs can be converted into CD4 Tregs restore IgE competence homeostatically via metabolic energy pathways. Indeed, a precedent was noted regarding a Qa-1-restricted natural formyl peptide of the nonclassical, nonpolymorphic MHCIb. This natural formyl peptide thus elicited universal protective CTLs against microbiota-induced skin disease in remarkably all strains of mice since Qa-1 as MHCIb class exhibits no polymorphism among all strains of mice ^51, 52^. Like Qa-1 in rodents, human HLA-E belonging to MHCIb class, consists of one single isoform without polymorphisms. Recently, human nonameric IgE peptides restricted to HLA-E was elucidated, which in principle can be universally applicable to human allergic patients ubiquitously inherited with this homozygous nonpolymorphic HLA-E ^13^. Thus, In view of the impending stride of universal IgE vaccines based on MHCIb vaccinology bypassing MHCIa polymorphism, a practical MHCIa CTL vaccines likewise must be inclusive by choosing appropriate MHCIa supertypes among the more than 19,000 alleles (as of May, 2020) ^53^. Henceforth in designing universal IgE allergic vaccines, nECPs-restricted to HLA-A2.01 and HLA-B7.02 together in the classical MHCIa herein encompass ∼48% of human population, which may be further expanded close to 99% by including three additional supertypes ^12^. Importantly, the identification of natural peptides through ER stress pathway, identifiable by the correspondence principle is of first-order relevance for designing vaccines for inflammatory diseases, autoimmune diseases and in particular viral infectious diseases ^54^. Herein we found induction of cell-mediated CD8 T cell cytotoxic immunity, that completely dampens IgE production. Notably, a naturally processed yellow fever viral peptide was identified exhibiting some tetramer-specific CD8 CTL clonotypes, however, unlike nECPs, this natural viral peptide did not offer pertinent CTL-mediated protection against yellow fever viral infection ^54^. In contrast, the endogenous immune-physiological nECPs characterized are major T cell epitopes the homeostatic metabolic third signal can transform the natural autoimmune self-peptide specific aggressor CD8+T cells that break natural tolerance into feedback CD4+ Treg.

### 4.2 The homeostatic Three-Signal Model: with DCs as Enabler/Generator (newly added)

#### 4.2.1 Historical milestones of associative recognition and S/NS discrimation

The emerging concept of one embodiment of tolerogenicity vs. immunogenicity stems from DW Dresser ^55^. In a milestone, the two signal theory of of PA Bretscher and M. Cohn (1970) ^56^ weds the concept of immune induction and immune tolerance into a theory of self/non-self (S/NS) discrimination. This qualitative framework of the immune system is distinct in principle from the quantitative model proposed by the carrier effect or association recognition model of Z. Ovary, B. Benaceraf, D. H. Katz, and A. Mitchinson ^57-59^. Using this model, nECP supertypes is a self-signal for emerging self-reactive precursor CTLs (pCTL), while promiscuous helper peptides (overcoming MHCII restriction) is the second signal that provide the CD4 helper signal. Semantically there is no self/non-self discrimination of an antigen so long the antigen is under a sufficient helper signal, enabled or licensed by costimulatory T cells and DCs, posing a protective sterile immunity on the one hand following induction to foreign antigens, and horror autotoxicus on the other hand, following breaking central tolerance to self-antigens. Tregs as a rescuer has been argued on an ad hoc base for regulating excessive anti-foreign or anti-self autoimmunity not yet conceptually integrated into a homeostatic antigen-specific S/NS/S framework ^60^.

#### 4.2.2 A mission critical extended conceptual framework for self integrity: S/NS/S and the GPT-Pex principle

Thus, although S/NS two signal theory unifies the quantitative associative recognition of foreign antigens in the costimulatory context of self antigens in making a qualitative distinction of self vs. non-self, the theory is incomplete for the lack of a basic principle for regaining self-tolerance and thus the protecting integrity of the self. Consequently, inductively activated cells following breaking tolerance rely heavily on senescence or metabolic decay of the diminishing second signals. Therefore, S/NS leaves the power of S/NS discrimination in a dangerous post hoc signaling zone, e.g., ‘The foundation is broken, the center cannot hold’. Therefore, there appears an urgency to un-reveal a mechanism of generative preexisting transformation (GPT) among the epitope presenting DCs, paratope-specific effector and regulatory cells, to back into self-tolerance in the three cell model using additional Tregs, sharing the same epitope/paratope-based minimal circuit.

In the current applied vaccinology, prolonged effector CD8 CTLs against the nECPs on IgE committed B cells and plasma cells risks deletion of IgE system resulting in the long-term IgE deficiency as observed in experimental perinatal IgE immunization via the breaking of IgE-class-restricted central tolerance in the laboratory ^1, 3-7^. As shown in Fig. 6, during homeostasis, CD8 nECP-specific CTLs can be peripherally transformed into the nECP-specific Treg, via generative DCs which in a metabolic milieu has unique activities to transform preexisting CTLs. Tregs transformed from CTLs remarkably lack the capacity in a division of labor to directly inhibit IgE production by IgE-producing cells, decorated with the same nECP on surface MHCI. Hence, to ensure IgE recovery. the transformed nECP-specific Tregs not only exhaust the source of CTLs, but also suppress the residual nECP-specific CTLs which escape transformation. The double edge sword of transformation is facilitated by the generative DCs in a versatile metabolic milieu (Fig 5).

**Fig. 6:**
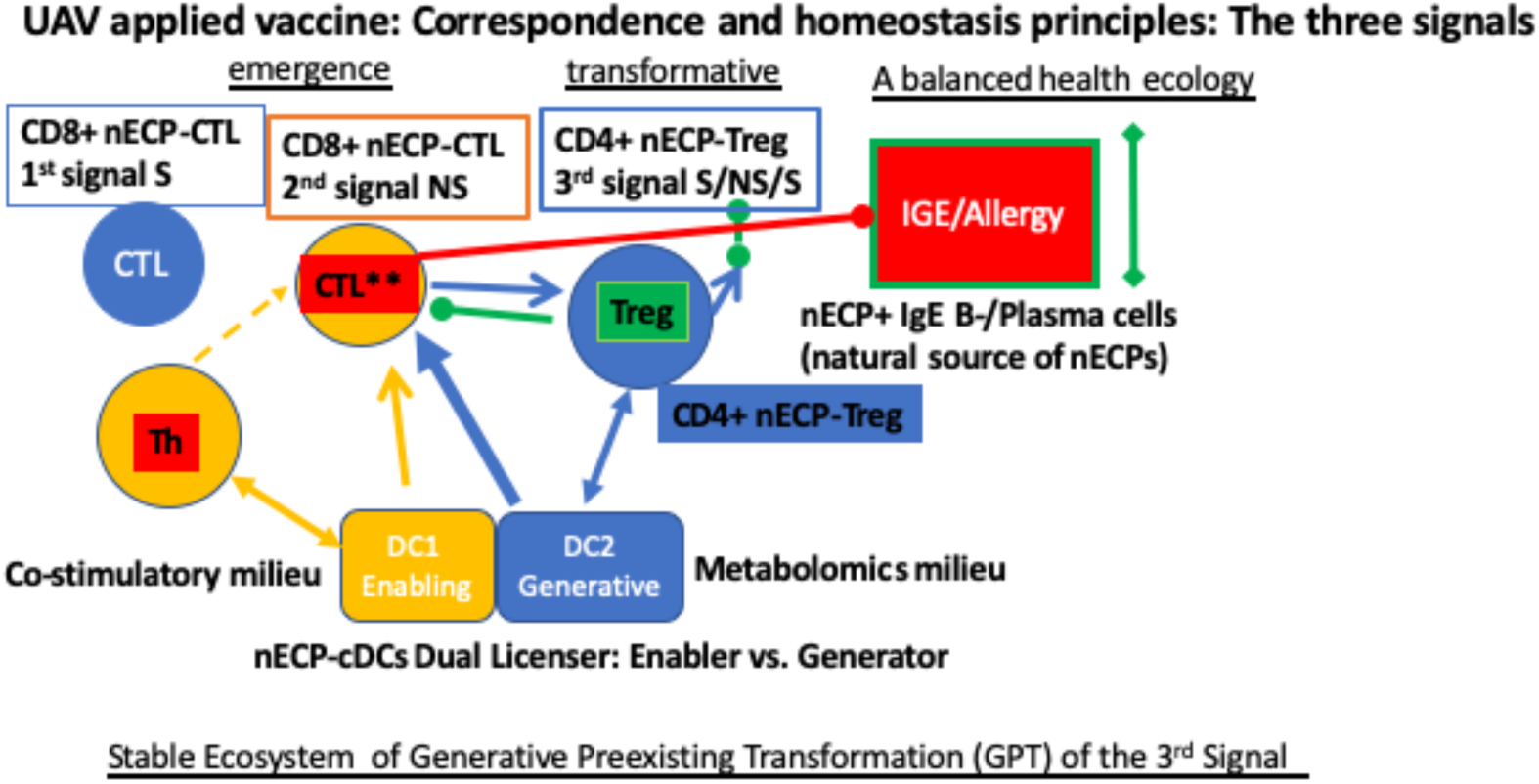
Three signal Generative-Preexistence-Transformation (GPT-Pex) for Self/Non-self/Self homeostasis. An homeostatic principle of nECP vaccinology: nECPs chemical entities as API the first signal for central self-tolerance (S); costimulatory second signal for inducing nECP- specific CTLs and breaking central tolerance (NS); and the transformation of preexisting CTLs into Tregs via generative DCs according to GPP-Pex, thus regaining peripheral self-tolerance (S) and the integrity of host IgE competence, following the removal of the IgE danger signal by mobile enabler DCs and CTLs in the dynamic three signal homeostatic S/NS/S.

Conspicuously, the nECP-based three signal model encompasses the homeostatic principle of regaining self-tolerance. The three signal concept is equivalent to the principle of the generative preexisting transformation (GPT-Pex), in which Generative DCs render the Preexisting effector be Transformed into Tregs in the closed minimal epitope-paratope circuit. No outsourcing is required for recruiting external Tregs. Hence GPT-Pex catapults the *bona fide* two signal theory of binary minimal S/NS formalism into a trinary minimum S/NS/S formalism. The three cell model marks three stage development of self-tolerance: Central tolerance of the emerging nECP-specific CTLs; NS: Breaking central tolerance by the second helper signal in eliminating IgE danger signal; S: Pre-existing effector CTLs be transformed into Tregs via nECP-generative DCs. The completion of the three stages regains homeostatic IgE immune competence.

Noticeably, the nECP tetramer-specific CD4+ Tregs converted from CD8+ CTLs maintained the expression of the cytokine biomarkers CD127/IFN-γ on CTLs, leaving the footprint of their origin (Fig. 1C). Interestingly, Tregs phenotype increase expressing T-bet at the population levels was found to correlate ad hoc with attenuated Th1-mediated colitis ^61, 62^ suggesting a GPT-Pex principle. The data of applied vaccination for the first time formally unite the second signal theory (S/NS) that can break loose to a three signal model S/NS/S in a minimum trinary circuit to keep aberrant autoimmunity at bay.

#### 4.2.2. Mechanisms of cell-cell interaction: DCs as Enabler vs. Generator

As shown in the bottom section of Fig. 6, DCs plays a dual role in S/NS/S. The three signal model comprises two sub-circuits pivoted on positive enabler DCs vs. negative generative DCs. During the phase of two signal breaking self-tolerance, DCs become the enabler for nECP-specific precursor CD8 (pCD8) herein for IgE natural peptides facilitated by helper peptides for CD4 T cells ^10^, or in a pivotal observation by Schoenberger and colleagues, CTLs for tumor antigen were licensed directly by anti-CD40 co-stimulated DCs ^63^. The role of a mobile DCs licensing anti-tumor immunity is importantly, formally demonstrated by Murphy and colleagues ^64^. In principle, the migratory capability of the class of enabler DCs vs. generative DCs increase efficacies of immune induction of effector T cells for protective immunity vs. licensing infectious Tregs to prevent horror autotoxicus.

In a physical three-cell constellation, targeting IgE-producing cells, pCD8 can be assisted by DCs in the proximity in a tripartite orientation via DCs, helper CD4 T cells and pCD8 T cells, since enabler DCs exhibiting nECP/MHCI as well as helper peptides/MHCII on the cell surface, relay respectively for CD8 and CD4 T cells during triparte interaction. Moreover, the enabler DCs, following the reciprocal help of helper CD4 T cells can in a different modality, directly enable or license pCD8 DCs. In similar veins, generative DCs under the metabolic influence can render transformation of CTL into Treg in the tripartite interactions, as well as directly license effector-Treg transformation. In contrast, direct T-T interaction appears unlikely since nECP is missing on MHCI of effector T cells, except frivolously piggybacking via the residual nECP vaccine; moreover, MHCII is absent on both MHCI+ CD4 and CD8 cells that annul MHCII-dependent helper or immunoregulatory interactions. Henceforth, the enabler vs. generative DCs, with increasing efficacies due to the mobility, underpin the relay and the dynamic transition of the three cells in a close physical interaction circuit shown in Fig. 6, which can be as many as seven different states including B cells/plasma cells as a physiological target.

In summary, the correspondence and homeostatic principles of the applied vaccinology may be both safe and efficacious. The nECPs-UAVs may treat categorically type I hypersensitivity of IgE-mediated allergies. Utility of the three signal vaccinology may be applied to ameliorate cytokine-mediated inflammatory diseases, and in particular, autoantigens-induced autoimmune diseases.

## Materials and Methods

### Human IgE antibody production

IgE producing cells cultures were initiated with PBMCs obtained from healthy donors selected by the SDBB, Inc. set registry according to self-reporting, wherein the IgE allergy status is not disclosed in the Agreement. PBMC contain multiple B cell lineages, including the IgE lineage and IgG lineage according to immunoglobulin production of the particular isotype, indicating the genetic switch of epsilon heavy chain gene rearrangements ^65, 66^ and commitment of IgE and IgG B cell lineage, e.g., IgE or IgG committed B cells and plasma cells, wherein the capacity of IgE and IgG production were selected in the study. Thus, 1x10^6^ human PBMC (single or preferably double, genotyped HLA-A2.01+/HLA-B7.02+ donors) were established as set up lineage IgE-producing cells using 7% H55 human plasma, anti-CD40 antibody (3 µg/mL) ^67^, IL-4 (300 U/well) ^65, 66^, IL-6 (60 ng/mL) ^68^, and IL-21 (60 ng/mL) ^69^. It is critical to obtain as many human plasmas and routinely screen for the capacity to support consistent human IgE production; with a competent carefully screened and selected plasma, IL-4 is still strictly required for IgE switch, while the cytokine supplementation, IL-6, IL-21 is dispensable. In our hand, all the PBMCs without exception thus far produce high levels of polyclonal IgE (up to 100 ng/ml or more) when cultured with appropriate human plasma. This remains the most important critical observation for enabling the analysis for regulating pathophysiological levels of human IgE production without skewly relying on human IgE producing SKO7 myeloma cells (HLA-A2.01/HLA-B7.02). Performing large batches of human donor plasma screening is rendered possible through the kind help and the generosity of Mr. Howard Brickner, at UCSD, San Diego, CA. The plasma screened for IgE production was treated with heparin with the removal of fibrinogen and clotting factors. This method has produced robust, consistent, and optimal batches for human IgE production for almost all the human PBMCs tested regardless of the diversity of the MHCI haplotypes.

Reagents and supplies and detailed methods are in details in the attached Supplement

## RESULTS

### I. Elucidating Naturally ER stress Processed Immunogenic or Tolerogenic Nonameric and Decameric Peptides

Analysis of binding to TAP-deficient T2 cells and induction of activated CD8 T cells by nonameric and decameric human IgE peptides

### [Extended Data] Table 1 to Table 3

PBMCs from San Diego Blood Bank were typed by primer sequencing to identify those expressing HLA-B7.02, HLA-A2.01 and HLA-A24.02, HLA-A3.01, HLA-A11.01, HLA- B51.01, HLA-B40.01 (Histogenetics, NY, NY). First, we explored binding of nonameric/decameric IgE peptides to HLA-A2.01 for upregulation of HLA-A2.01 on TAP-deficient T2 cells that express the endogenous HLA-A2.01. A panel of IgE peptides (code #17 to #50) in Table 1A with the predicted high binding affinity to HLA-A2.01 according to the IEBD MHCPan4.0 program (LJI, La Jola, CA) were prepared as synthetic peptides. Notably, none of these synthetic peptides are capable of binding and upregulating surface HLA-A2.01 of the TAP-deficient T2 cells (not shown). Therefore, we next employed an alternative direct CD8 T cell activation assay using the method pioneered by Appay and colleagues ^1^ for screening the HLA-A2.01 synthetic peptides (code A21 to A36) in Table 1B IgE nonameric or decameric peptides as listed in Table 1B predicted according to the BIMAS program (NIH, Bethesda, MD). Notably, none but the A32 peptide exhibited strong induction of CD8 T cells. Thus, A32 was shown to induce CD8 T cells in HLA-A2.01 expressing PBMCs concomitantly stimulated with promiscuous PADRE (AKAVAAWTLKAAA) or <u>M</u>easles <u>h</u>elper <u>p</u>eptides (MHP, ISISEIKGVIVHKIETILF) for cytokine activated CD8 T cells (listed in Table 1B).

**Table 1A.**
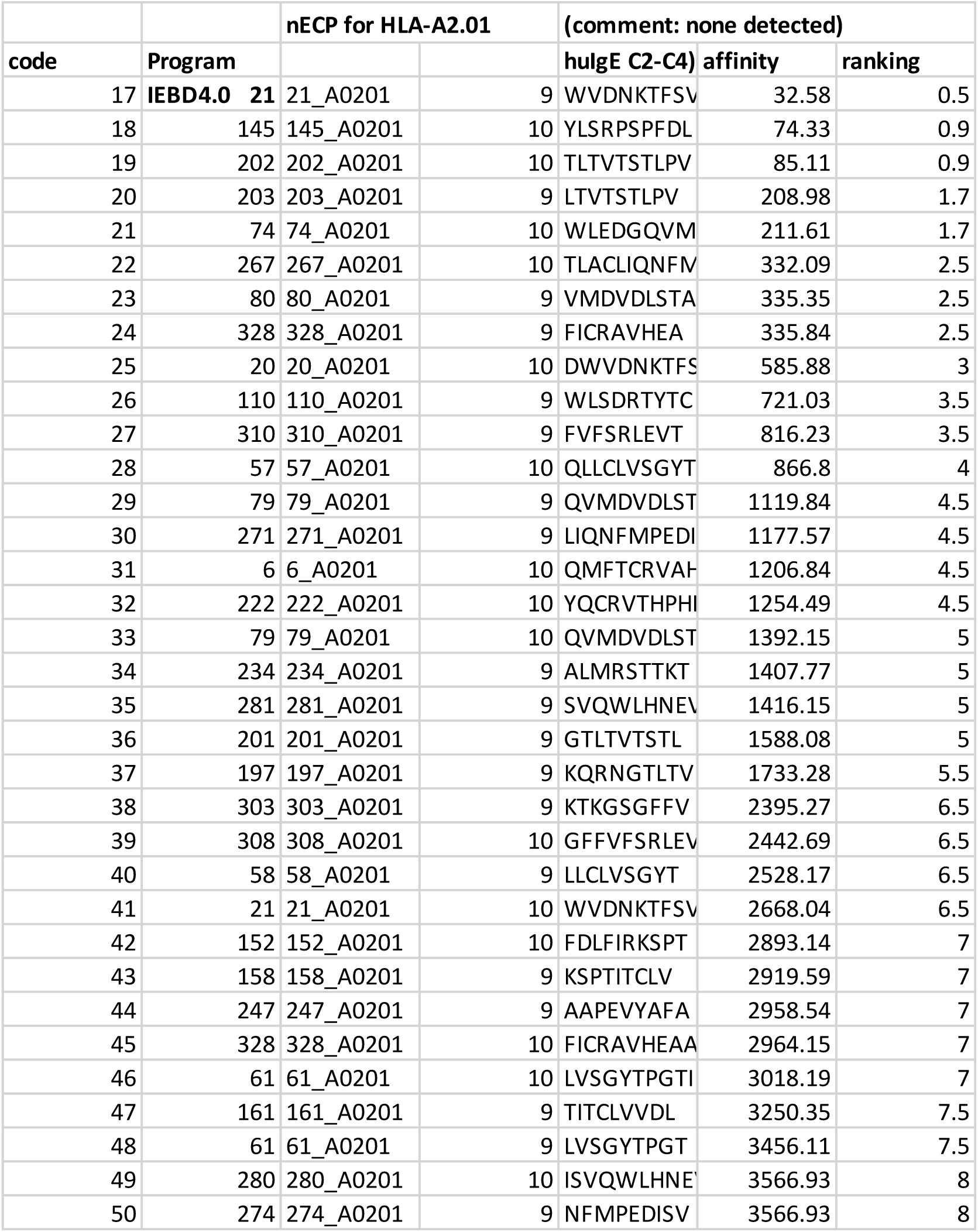

**Table 1B.**
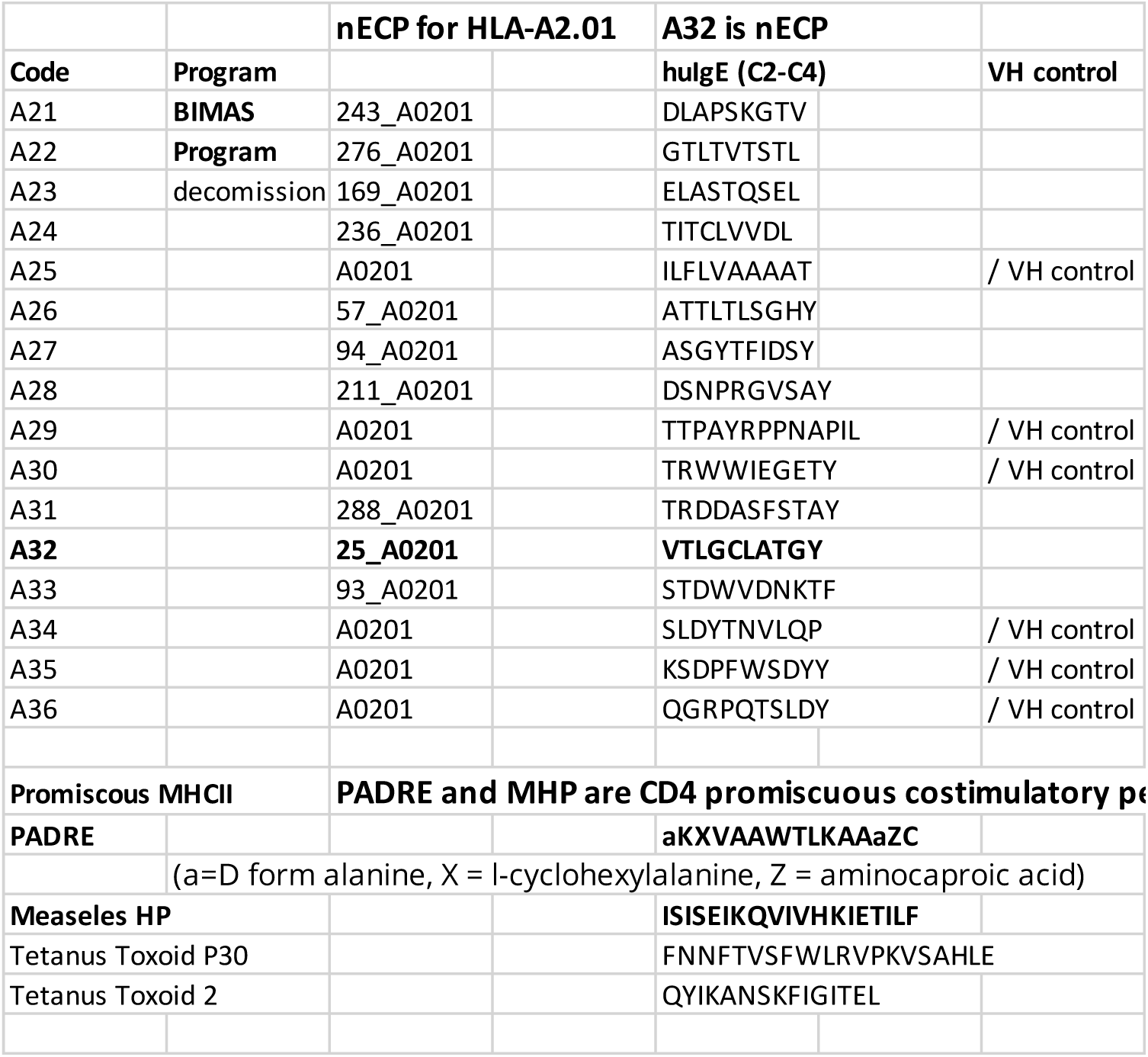

Next, we explored binding of nonameric/decameric IgE peptides binding to HLA-B7.02. A TAP-deficient HLA-A2.01 expressing T2 cell line, stably transfected with the HLA- B7.02 gene was kindly obtained from Professor Charles Lutz at the University of Kentucky) ^2^. IgE synthetic peptides (Code #153 to #186) were prepared in Table 2 according to B7.02 binding motif of the IEBD NetMHCpan-4.0 Program, and were tested for upregulating surface HLA-B7.02 expression in the B7.02 gene transfected TAP deficient T2 cell line. Notably, IgE nonamer, SP1 (HPHBPRALM), SP2 (NPRGVSAYL) as well as the positive control nonameric peptide of EBV B7 (RPPIFIRRL) upregulated surface HLA-B7.02 as detected by the anti-HLA-B7.02 fluorescent antibody set. Next, we further extended the detection of IgE synthetic peptides for eight different MHCI supertype loci according to Table 1C. Since the respective MHCI supertype-gene transfected TAP-deficient T cell line was not available, we therefore employed the direct T cell activation assay for screening potential MHCI binding peptides for the six different HLA-A and HLA-B supertypes using the method of Appay ^1, 3^. Notably none of the prepared synthetic peptides induced CD8 T cell activation as compared to that induced by A32 nonameric peptide restricted to HLA-A2.01 as a positive control.

**Table 2.**
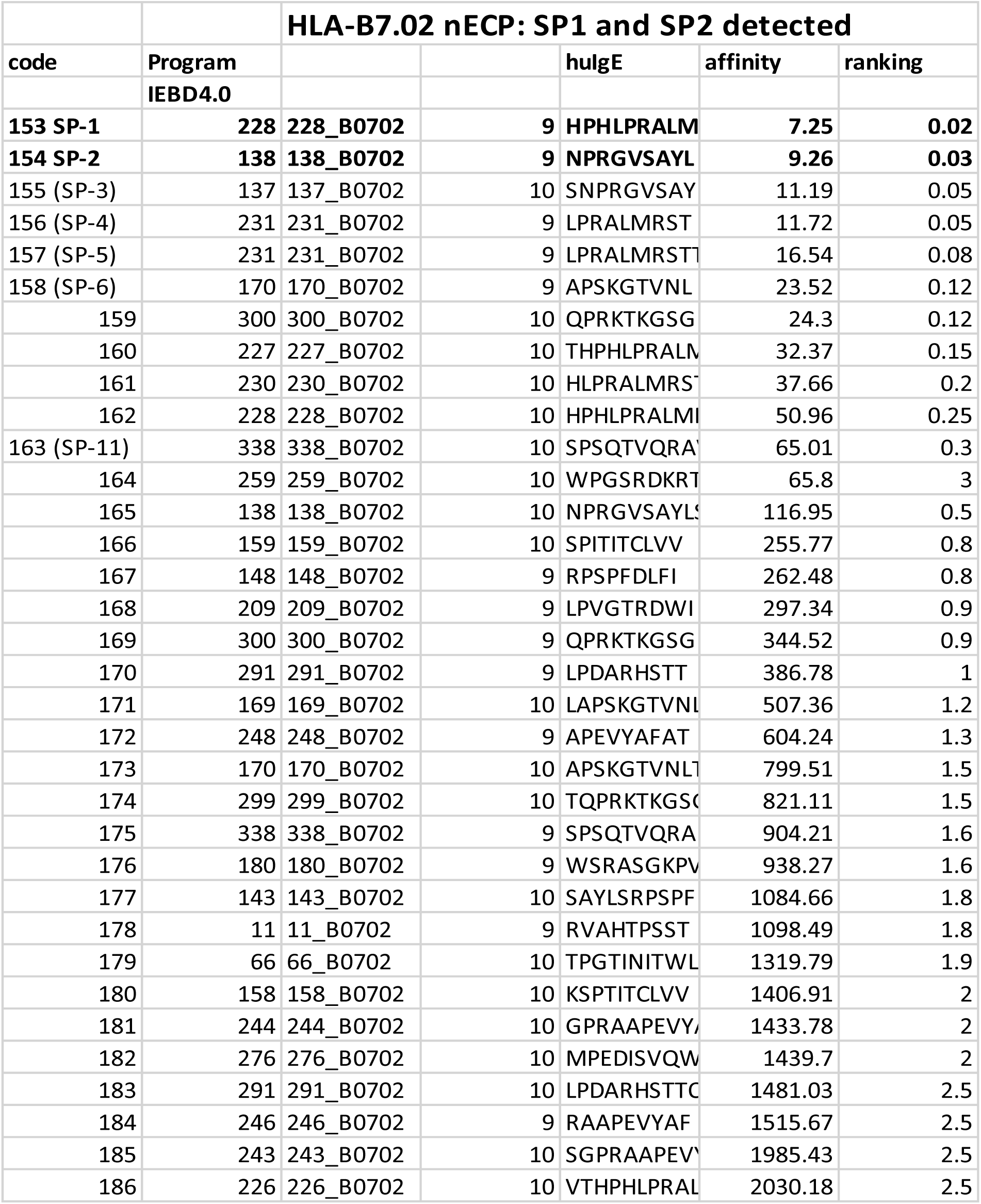

### [Extended Data Suppl.] Table 1 to Table 3 IgE synthetic peptide binding to human MHCI supertypes via surface MHCI upregulation vs. direct T cell activation

### II. Construction of Triple Transgenic Animal Model: Human HLA- A2.01 and HLA-B7.02 Restricted Transgenic Human IgE as Antigens Expressing on DCs and Monocytes of the Triple Transgenic animals

**[Extended Data Suppl.] Fig. 1.**
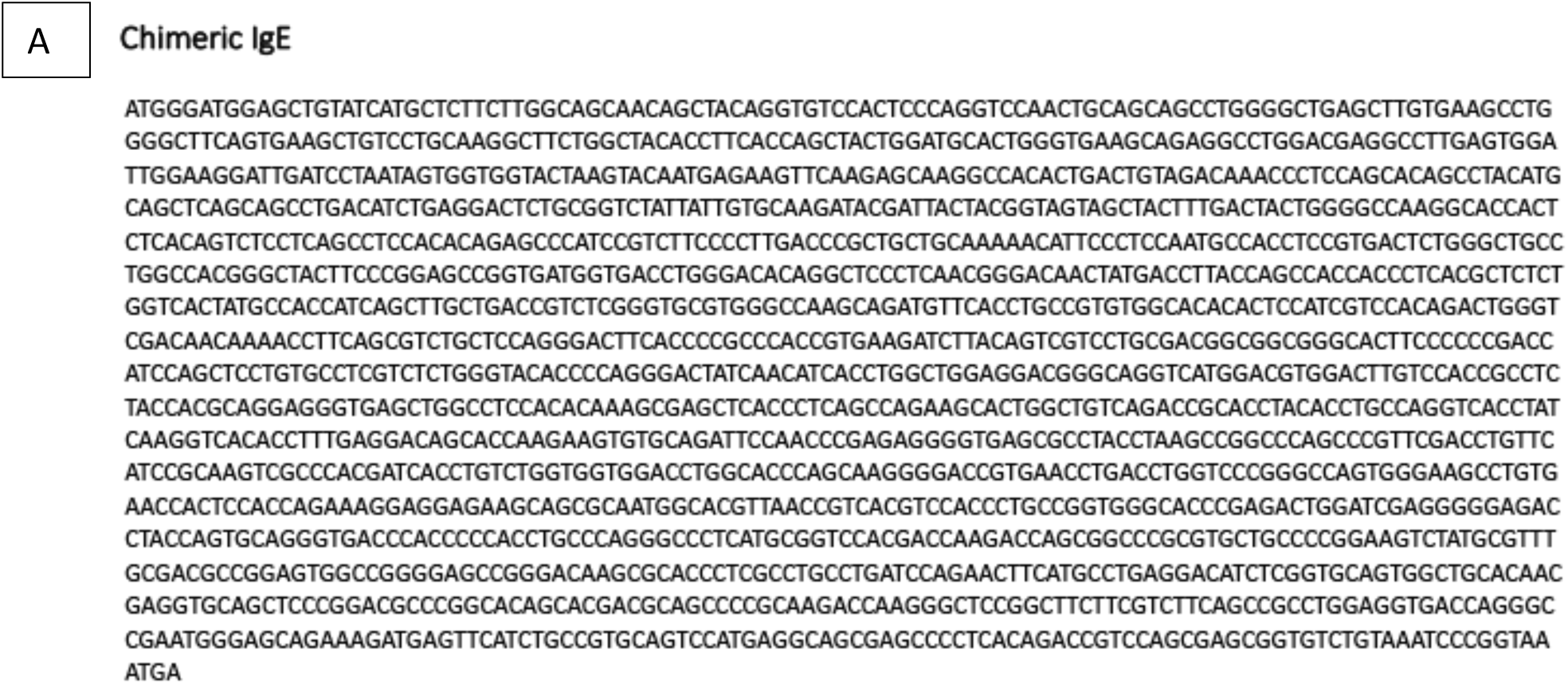
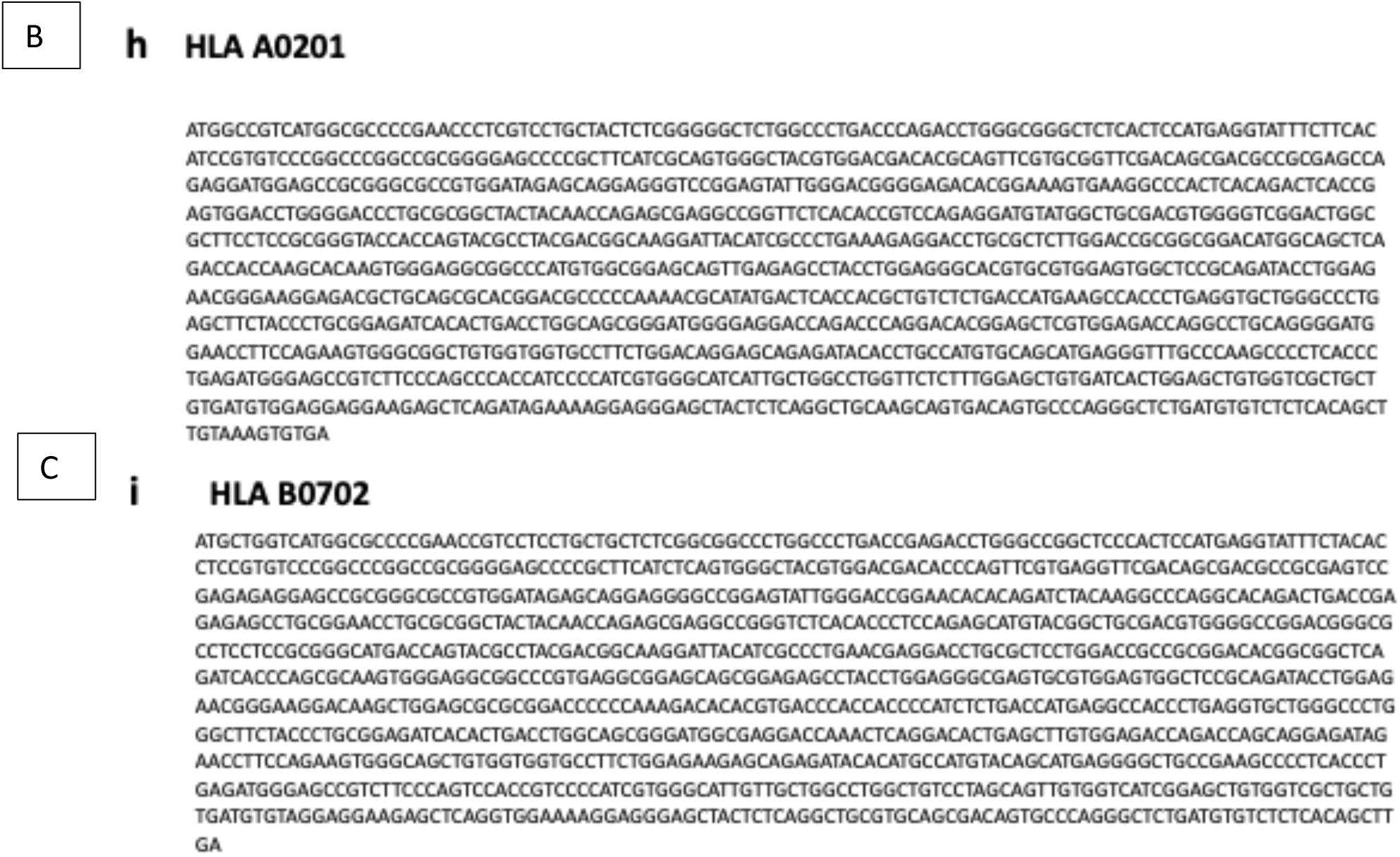
Construction of triple transgenic animals. A colinear DNA of triple human transgenes of HLA-A2.01, HLA-B7.02, and chimeric JW8 human epsilon heavy chain, under the respective CMV and EF1a promoter control, was injected into the pronucleus of fertilized eggs to produce triple human transgenic mice. Mouse strain: C57BL/6x C57BL/6 (DOB: 01-21-2019), with 2 female and 2 female pups as founder #30 and #39; and two male selected founder #26 and #36 out of 51 injected pups, screened with the PCR gene probes. Genomic DNAs were prepared from the mouse tails. IgE, HLA-A02.01, and HLA-B07.02 transgenes were detected by PCR with the respective primer sets. DNA sequences of the chimeric JW8 IgE transgene, HLA-2.01 and HLA-B7.02 transgenes were shown (Suppl. Fig. 1A-C), The mice were backcrossed for nine generations to the parental C57BL/6, and the genomic triple transgenes were typed by PCR. The concomitant and colinear mRNA expression of the transcribed triple genes was shown by RT-PCR.

**[Extended Data Suppl.] Fig. 2.**
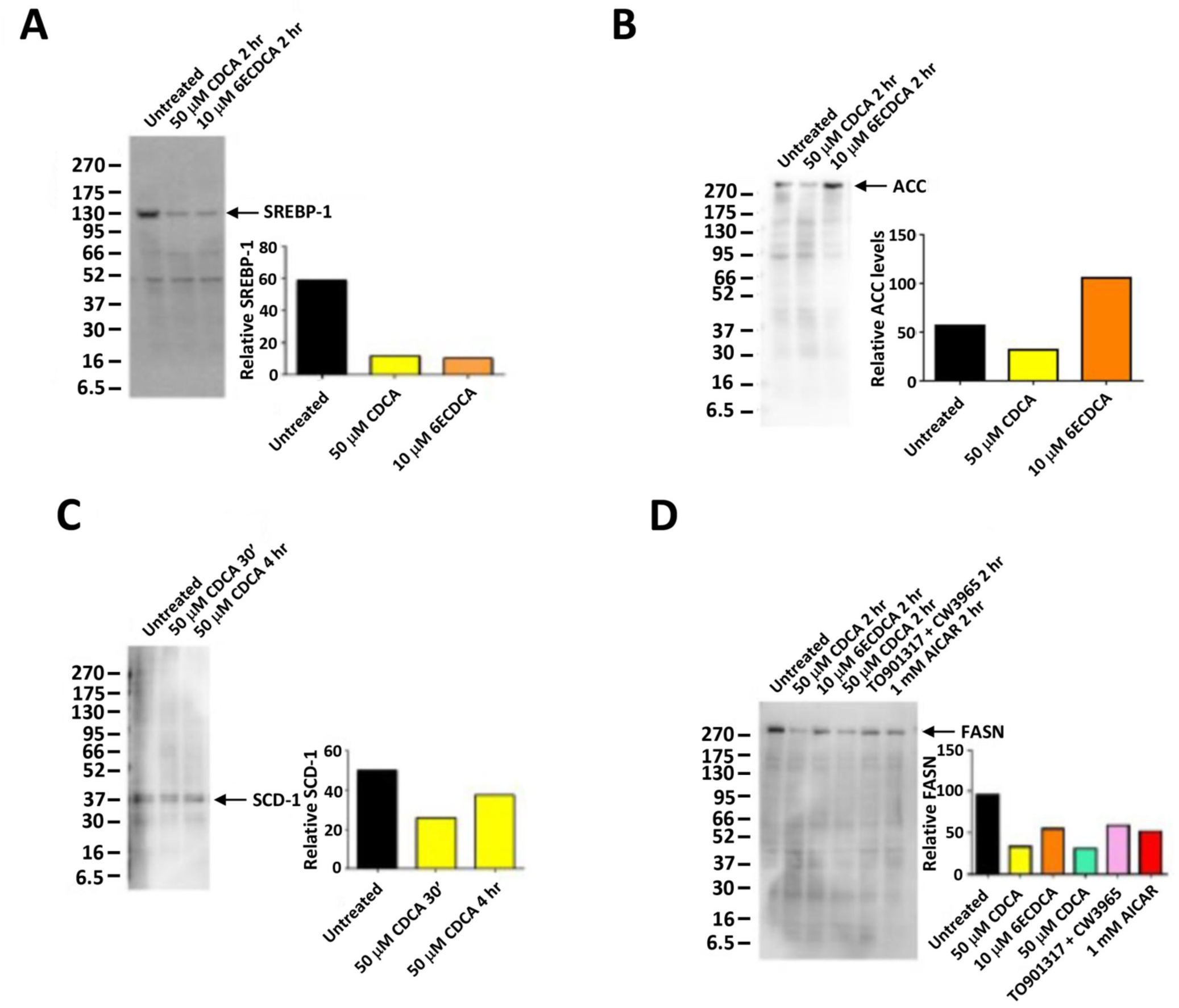
Regulation of SREBP-1c and the downstream lipogenic genes in the HepG2 cell line. HepG2, a hepatocellular carcinoma, is widely used for studying lipid metabolism as a model for obesity. HepG2 constitutively expresses high levels of master transcription factor SREBP-1c, which upregulates lipogenic genes: acetyl-CoA carboxylase (ACC1/2), stearoyl CoA desaturase (SCD1), and fatty acid synthase (FAS/FASN) 4. ACC catalyzes malonyl CoA, the building block of long-chain fatty acids; SCD1 plays the role of unsaturated fatty acid synthesis for oleic acid, while FAS captures acetyl-CoA/malonyl CoA at the N-terminus, donating to the C terminus intermediate chain to assemble into C-16 palmitic long-chain fatty acid as fuels. Here we showed (Suppl. Fig 4 A-D) that SREBP-1c was downregulated 88% by the canonical CDCA and 6E-CDCA (A); ACC1/2 was downregulated by CDCA, however, upregulated by 6E-CDCA (B); and SCD1 was downregulated by 31% to 60% by CDCA or 6E-CDCA treatment (C). Moreover, FASN downstream of SREBP-1c was downregulated ∼73% by CDCA, 48% by 6E-CDCA, 75% by OEA, and 49% by AICAR/metformin (D). Thus, in addition to the liver cells, typically used for studying energy metabolism, it is important also to investigate the differential use of the SREBP-1c pathway for MoDCs governing transition of CD8+ CTLs vs. CD4+ Tregs.

### Legend to [Extended Data] Table 1 to Table 3

a. TAP assay: TAP-deficient T2 cell line with endogenous HLA-A2.01 is used for screening HLA-A2.01 binding synthetic IgE peptides in Table 1A for HLA-A2.01 binding (according to the IEBD NetMHCpan-4.0 program) for cell surface upregulation at nonpermissive temperature at 37°C as well as synthetic IgE peptides in Table 1B for HLA-A2.01 binding (according to the NIH BIMAS program). Another TAP-deficient T2 cell line stably transfected with the HLA-B7.02 gene was used for screening HLA-B7.02 IgE synthetic peptides according to (according to the IEBD NetMHCpan-4.0 program). The HLA-B7.02- transfected cell line, kindly provided by Professor Lutz at the University of Kentucky *^2^* was maintained in 300 μg/ml hygromycin. T2 B7.02 cells were pulsed with the different peptides (10 μg/ml) listed in Table 2 overnight at the nonpermissive temperature 37°C, wherein the non-binding peptides will not permit surface HLA-B7.02 expression at 37°C, and then stained with an anti-HLA-B7.02 antibody (BioLegend, Cat: 372402) and an Alexa Fluor 488-conjugated anti-mouse antibody for FACS evaluation.

b. CD8 T cell activation assay (TAA) of the method of Appay ^1, 3^. PBMCs from San Diego Blood Bank (SDBB) were typed by primer sequencing to identify those expressing HLA-B7.02, HLA-A2.01 and HLA-A24.02, HLA-A3.01, HLA-A11.01, HLA- B51.01, HLA-B40.01 (HistoGenetics, NY, NY), which were screened directly through synthetic IgE peptides according to IEBD and NetMHCpan-4.0 for cytokine-activated CD8 T cells, co-stimulated with promiscuous PADRE (AKAVAAWTLKAAA) or measles helper peptide (MHP) (ISISEIKGVIVHKIETILF) (Table 1B, Table 3 and Table 4). Using the method of Appay ^11^, PBMCs were incubated at 5 x 10^5^ cells/well in a 96-well plate in AIM-V medium (Gibco) supplemented with 50 ng/ml of FLT3L and plated. The next day (day 1), 0.5 µg/ml of ssRNA40,10 µg/ml of A32 peptide (VTLGCLATGY) 2 µg/ml of PADRE helper peptide ^1, 3^ (screened from IgE peptides from the constant IgE heavy chain (Cε1-Cε4) vs. control peptides (encoded by the VH region candidates) were added, compared to the positive control Melan-A/MART-1 peptide (EAAGIGILTV, not shown) or ELA-10 octameric peptides (ELAGIGILTV). For HLA-A2.01 screening purpose, on day 10, mitomycin C-treated (25 µg/ml) HLA-A2^+^ EBV-transformed B-cells, A32 (10 µg/ml) were added to wells in the presence of 1 µg/ml of anti-CD28 and anti-CD49d antibody costimulation and then incubated for 6 h at 37°C. Un-pulsed EBV-infected B-cells were added in parallel as a control for nonspecific background. For the final 5 h of incubation, 2.5 µg/ml of monensin and 5 µg/ml of brefeldin A were added. Cells were then collected and stained Alexa 488-conjugated anti-CD8 and PerCP/Cy5.5-conjugated anti-CD137 antibodies (extracellular) and APC-conjugated anti-IFN-γ and pacific blue-conjugated anti-granzyme B antibodies (intracellular). Flow cytometry was conducted with an LSR-II analyzer and FlowJo software.

**Table 3.**
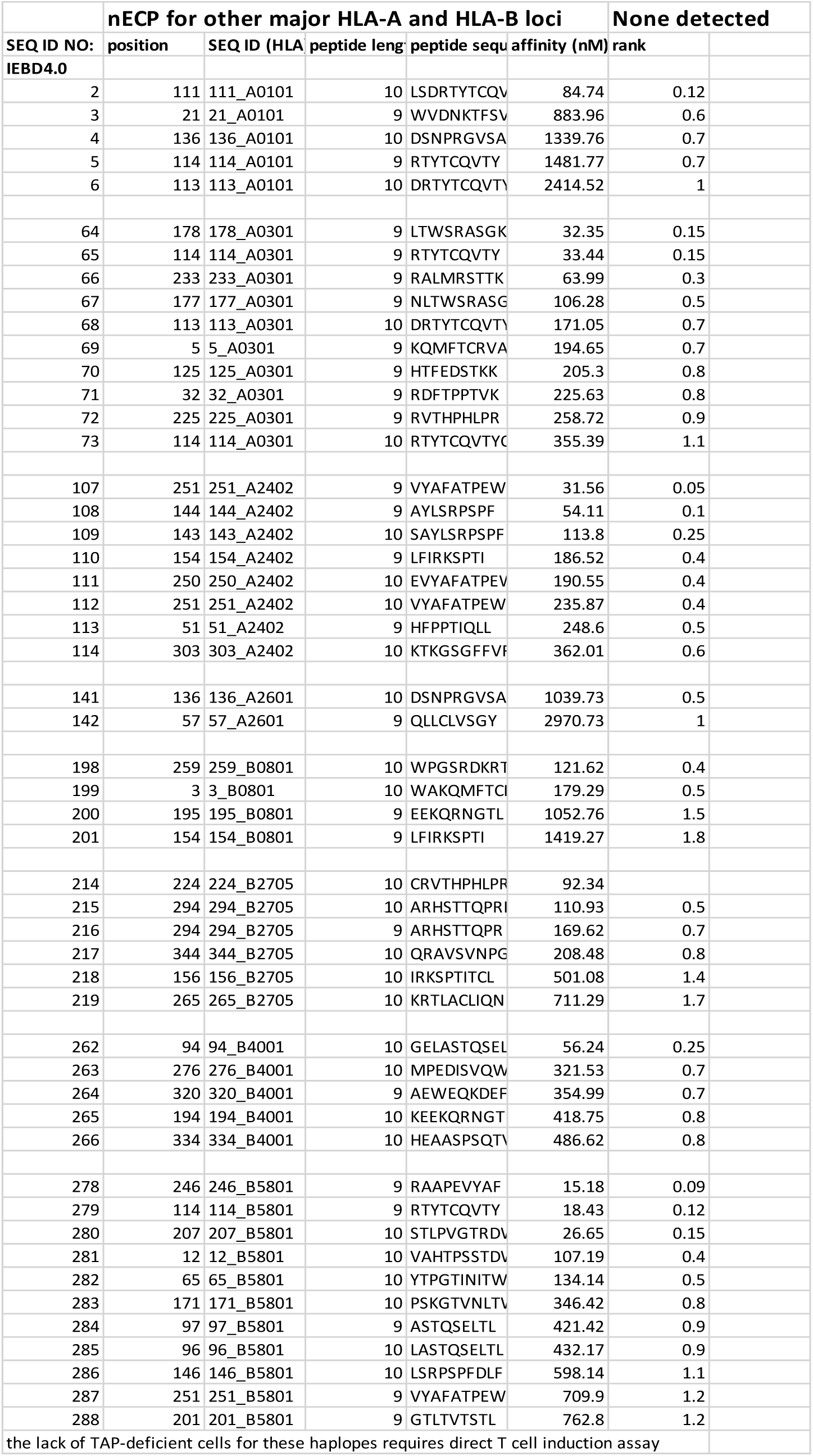

### Legend to [Suppl.] Fig. 1 Construct of triple transgenic mice expressing human HLA-A2.01, HLA-B7.02 and chimeric JW8 human IgE

Colinear construct for chimeric-IgE transgenic mice. HLA A0201, HLA B0702, and IgE under the respective control of CMV and EF1a promoters, was injected into the pronucleus of fertilized eggs to produce the founder transgenic progeny. Genomic DNA was prepared from mouse tails using a genomic DNA preparation kit. IgE, HLA A0201, and HLA B0702 were detected by PCR with primers CTTCCATTTCAGGTGTCGTG (forward) and CAAGCTGA TGGTGGCA TAGT (reverse) for IgE, CACGTAGCCCA CTGCGATG (forward) and CAATGGGAGTTTGTTTTGGCACC (reverse) for HLA- A02.01, and CTACGACGGCAAGGATTACAT (forward) and GGGCAGCTGTGGTGGTGC (reverse) for HLA-B07.02. The PCR conditions were 35 cycles of 94^0^C for 20 s, 60^0^C for 30 s, and 72^0^C for 35 s. **c.** RT-PCR was used to assess the mRNA expression of IgE, HLA-A0201, and HLA-B0702. Cells were collected from mouse blood after depletion of red blood cells using RBC lysis buffer, followed by RNA purification with an RNA purification kit. The cDNAs were obtained by reverse transcriptase (Bio-Rad) and the following primers were used: GCCACACTGACTGTAGACAAA (forward) and CAAGCTGATGGTGGCATAGT (reverse) for IgE, TGGACAGGAGCAGA GATACA (forward) and ACAAGCTGTGAGAGACACATC (reverse) for HLA-A0201, and CTACGACGGCAAGGATTACAT (forward) and AGCTCAGTGTCCTGAGTTTG (reverse) for HLA-B07.02. Primers CAGGAGAGTGTTTCCTCGTCC (forward) and GATGGGCTTCCCGTTGATGA (reverse) were used for the GAPDH internal control.

### Legend to [Suppl.] Fig 2. Effect of drugs on SREBP-1 gene and downstream lipogenic genes by HepG2 cells

Human HepG2 cells [# HB-8065, American Type Culture Collection (ATCC), Manassas, VA] were maintained in Eagle’s Minimum Essential Medium (EMEM) medium (ATCC # 30-2003), critically important for its survival and cannot be substituted by other types of media, containing 20% FBS at 37°C with 5% CO2. For drug treatment in the same medium, 2 ml of cells (2 x 10^6^) cells per 3.5-cm dish were cultured overnight and then treated as indicated. For drug treatment under the serum starvation condition, HepG2 cells (2 x 10^6^ cells per 3.5-cm dish) were cultured in serum-free EMEM medium (1 ml per 3.5-cm dish) for 48 hours followed by drug treatment as indicated. After drug treatment, cells were lysed by CHAPS lysis buffer, and for western blotting, 15 μl of cell lysate (15-30 μg of proteins) per lane were resolved by 4-20% SDS-PAGE as described in the METHODS. (a) The SREBP-1c: Primary antibody: mouse anti-SREBP-1 monoclonal antibody (# sc-365513, Santa Cruz Biotechnology, Dallas, TX; 1:200). Secondary antibody: HRP-conjugated goat anti-mouse IgG (H+L) secondary antibody (# 32430, Thermo Fisher Scientific, Waltham, MA; 1:1000). (b) ACC1/2: Primary antibody: rabbit anti-ACC monoclonal antibody (# 3676T, Cell Signaling Technology, Danvers, MA; 1:1000). Secondary antibody: HRP-conjugated donkey anti-rabbit IgG (#406401, BioLegend, San Diego, CA; 1:1000). (c) SCD-1: Primary antibody: sheep anti-human SCD-1 antibody (# AF7550-SP, R&D Systems. Minneapolis, MN; 1:2000). Secondary antibody: rabbit anti-sheep IgG (H+L)-HRP (#6150-05, SouthernBiotech, Birmingham, AL; 1:1000). (d) FAS/FASN: Primary antibody: rabbit anti-FASN antibody (# 3189S, Cell Signaling Technology, Danvers, MA; 1:1000). Secondary antibody: HRP-conjugated donkey anti-rabbit IgG.

#### Source of human peripheral blood lymphocytes (PBMCs)

PBMC from more than 50 healthy donors from the San Diego Blood Bank, are PCR genotyped by genotyping service in NYC for HLA-A and HLA-B alleles, from whom ∼22 individuals to be double or single positive for HLA-A2.01, and/or HLA-B7.02. were used throughout the coculture experiments. The male and female are ∼70% vs. 30%, with Caucasians (∼40%; Hispanics, 35%, and Asia and Pacific Islanders ∼25%. The medical status of healthy donors including IgE allergy status are confidential and released in the registry of SDBB. The usage of human volunteer blood donors is approved under license of IRB for IGE Therapeutics and the Institute of Genetics: IORG 0008xxx and IgE &Genetics IRB#x-Xxxrgy. The plasma are collected from the individuals. If the PBMCs from the donors whole blood, the buffy coats are prepared and typed immediately by FACS for surface HLA-A2 (mAb BB7.2) and HLA-B7 (mAb BB7.1) by mAbs from BioLegend, Inc., and later the HLA-A2.01 and HLA-B7.02 further confirmed with PCR genotyping (Histogenetics, N.Y.). Data will not identify a donor, unless (a) specifically provided for in the Agreement, and (b) permitted or required by law (including, without limitation, Donor consent or authorization and/or IRB (as that term is defined below approval). The Agreement specifies the Purpose for use of the materials, the donor data, and any test results provided under the purpose of use of the Agreement. The Purpose of the Agreement is for the research purpose to advance biomedical knowledge advancement and data publication in peer-reviewed journal as well as scientific conference or local functions for educative purpose, and not selected for purpose in “Treatment of a human being” in accordance with standard medical practice and the FDA regulations and guidelines, excluding requirements for investigational new drug (IND) applications, drug, biological products or medical devices regulations, human cells, tissues, and cellular and tissue-based products standards, and current good manufacturing and tissue practices, and other Applicable Law.

#### Reagents

Complete RPMI 1640 medium (10% FBS, 1% L-Glutamine and 1% Penicillin/Streptomycin), 10-cm cell culture dish (Olympus plastics; Cat: 25-202), recombinant human FLT3L (BioLegend Cat: 550606); IL-4 (Stock: 600 units/µL; Final: 200 units/ml), BioLegend; Cat: 574006, GM-CSF (Stock: 20,000 units/µL; Final: 100 units/ml), IL-1β (Stock: 200 μg/ml; Final: 5 ng/ml), TNF-α (Stock: 200 μg/mL; Final: 10 ng/ml), IL-6 (Stock: 200 μg/ml; Final: 60 ng/ml), BioLegend; Cat: 570804, Prostaglandin E2 (PGE2) (Stock: 2.5 mg/ml; Final: 1 μg/ml), 96-well flat bottom culture plate (Olympus plastics; Cat: 25-109), OEA, 5 ml (12 x 75 mm) polystyrene round bottom FACS tubes, FACS Staining Buffer (2% FBS in DPBS), DMEM medium (GenClone; Cat: 25-500N) containing 7% H55 human plasma, 1% L-Glutamine and 1% Penicillin/Streptomycin, LEAF^TM^ Purified anti-human CD40 antibody, 3 μg/ml (BioLegend; Cat: 334304; 1 mg/ml), IL-4 (300 U/well) (BioLegend; Cat: 574004; 0.2 mg/mL, 0.5-2.5 x 10^7^ units/mg, 1,000 U/μl), IL-6 (60 ng/mL), IL-21 (60 ng/ml) (BioLegend; Cat: 571202; 0.2 mg/ml), LEAF Purified anti-human NKG2D (CD314) antibody (BioLegend; Cat: 320809), LEAF Purified anti-human MICA/MICB antibody (BioLegend; Cat: 320909), Purified anti-human CD94 antibody (BioLegend; Cat: 305502), Human Trustain FcX (Fc Receptor Blocking Solution: BioLegend; Cat: 422302), FOXP3/Transcription Factor Staining Buffer Set (eBioscience; Cat: 00-5523), Brefeldin A (10 μg/ml, BioLegend; Cat: 420601), BV570 anti-human CD4 antibody (BioLegend; Cat:300534), AF700 anti-human CD8 antibody (BioLegend; Cat: 344724), AF488 anti-human FOXP3 antibody (BioLegend; Cat: 320012), BV421 anti-human TGF-β antibody (BioLegend; Cat: 349613), APC/Cy7 anti-human CD25 antibody (BioLegend; Cat: 302614), APC anti-human IFN-γ antibody (BioLegend; Cat: 502512), PerCP/Cy5.5 anti-human CD137 (4-1BB) antibody (BioLegend; Cat: 309814), PE/Cy7 anti-human IL-10 antibody (BioLegend; Cat: 501419), human IgG4 (BioLegend; Cat: 403701) All small molecules were purchased from Tocris, Inc. (Minneapolis, MN) or Cayman Chemical (Ann Arbor, MI): AMPK mix (AMPKm: 1 mM AICAR, 100 nM Metformin), calcitriol (VD3; 100 nM), bexarotene (Bexa; 100 nM), SR11237(100 nM), or retinoic acid/vitamin A (100 nM), or GSK-J4 (30 mM), Oleoylethanolamide (OEA, 20 mM), PPARa agonist, WY-14643 (20 mM). All the peptides were synthesized by Biomatik, Inc. (Ontario, Canada), EBV B7 (RPPIFIRRL), IgE nECP nonamers, SP1 (HPHBPRALM), SP2 (NPRGVSAYL), IgE nECP decameric A32 peptide (VTLGCLATGY), PADRE (AKAVAAWTLKAAA) or measles helper peptides (MHP, ISISEIKGVIVHKIETILF). Tetramers of the nonameric PE-SP1_HLA-B7.02 and PE-SP2_HLA-B7.02 were prepared by the NIH Tetramer Core Facility (Atlanta, GE), while the decameric PE-A32_HLA2.01 failed after repeated attempt. Human Hep G2 cells [# HB-8065, American Type Culture Collection (ATCC), Manassas, VA]; Eagle’s Minimum Essential Medium (EMEM) medium (ATCC # 30-2003). Human peripheral blood mononuclear cells (PBMCs) were treated with GM-CSF and IL-4 to induce monocyte derived dendritic cells (MoDC). Cells were further treated with IL-1β, IL-6, TNFa and prostaglandin E2 (PGE2) for maturation.PGE-2/IL-6/IL-1β stimulated mature MoDCs were treated with different metabolic modulators or siRNA at the indicated doses and time courses. Cell lysates were analyzed by Western blot using anti-SREBP- 1c (sc-365513); anti-FASN (cst #3189S); anti-SHP (# ab96605); anti-CTLA-4 (BNI3). Levels were quantified using ImageJ software. Knockdown of SREBP-1c by siRNA correlated with FOXP3 upregulation by qRT-PCR.

## METHODS

### I. Elucidating the Immunogenic Protective DCs and MoDCs Nonameric and Decameric Peptides

The three-step method: binding, induction, and verification to fulfill the Correspondence Principle

### Step one: Peptide binding to TAP1/2 deficient cell lines

The binding assay is used for determining potentiality of the peptides as vaccine with putative high affinity binding to upregulate surface MHCI. TAP-deficient T2 cells transfected with the HLA-B7.02 gene (Dr. Lutz, University of Kentucky) ^2^ and maintained in 300 μg/ml hygromycin. T2 B7.02 cells were pulsed with the different peptides (10 μg/ml) listed in Table 1A overnight in nonpermissive temperature 37°C, wherein the non-binding peptide will not permit surface HLA-B7.02 expression, and then stained with an anti-HLA-B7.02 antibody (BioLegend, Cat: 372402) and an Alexa Fluor 488-conjugated anti-mouse antibody for FACS evaluation.

### Step two: CD8 T cell activation (TAA) assay

the method of Appay determines the vaccine potentiality when the peptides exhibited no activities to upregulate surface MHCI ^1^. PBMCs from the Blood Bank from healthy donors were typed by primer sequencing for HLA-B7.02, HLA-A2.01, HLA-A24.02, HLA-A3.01, HLA-A11.01, HLA-B51.01, HLA-B40.01 (Histogenetics, NY, NY), which were screened directly through synthetic IgE peptides according to IEBD and MHCPan4.0, and BIMAS for cytokine-activated CD8 T cells, co-stimulated with promiscuous PADRE (AKAVAAWTLKAAA) or MHP (ISISEIKGVIVHKIETILF). PBMCs were incubated at 5 x 10^5^ cells/well in a 96-well plate in AIM-V medium (Gibco) supplemented with 50 ng/ml of FLT3L and plated. The next day (day 1), 0.5 µg/ml of ssRNA40,10 µg/ml of synthetic peptides and 2 µg/ml of PADRE or MHP helper peptide ^1, 3^ (screened from IgE peptides from the constant IgE heavy chain (Cε1-Cε4) vs. control peptides VH region candidates) were added, compared to the positive control Melan-A/MART-1 peptide (EAAGIGILTV) or ELA-10 peptides (ELAGIGILTV). For HLA-A2.01 screening purpose, on day 10, mitomycin C-treated (25 µg/ml) HLA-A2^+^ EBV-infected B-cells pulsed with A32 (10 µg/ml) were added to wells in the presence of 1 µg/ml of anti-CD28 and anti-CD49d antibody costimulation and then incubated for 6 h at 37°C. Un-pulsed EBV-infected B-cells were added in parallel as a control for nonspecific background. For the final 5 h of incubation, 2.5 µg/ml of monensin and 5 µg/ml of brefeldin A were added. Cells were then collected and stained Alexa 488-conjugated anti-CD8 and PerCP/Cy5.5-conjugated anti-CD137 antibodies (extracellular) and APC-conjugated anti-IFN-γ and Pacific blue-conjugated anti-granzyme B antibodies (intracellular). Flow cytometry was conducted with an LSR-II analyzer and FlowJo software.

### Step three: Verification of the Correspondence Principle, the actuality of the nECP vaccine

To further test whether the activated CD8 T cells inhibited IgE production by IgE-producing B cells, PBMCs cultured as above were stimulated with SP-1, SP2, or A32 in the presence of PADRE or MHP (10 μg/ml) ^5, 6^, treated with or without the above AMPK mix (1 mM AICAR, 100 µM metformin), VB, OEA, or GSK-J4. On day 7 to day 10, the cells were harvested, and incubated with peptide pulsed T-2 indicator cells for lysis and IgE producing cells (i) CTLs from vaccinated PBMCs were then incubated with A32, or SP-1/SP-2 pulsed HLA-B7.02 transfected T2 cells vs. non-peptide-pulsed T2 cells in an LDH assay (Thermo Fisher kits): Briefly, 50 μl of supernatant and 50 µl of substrate mixture were incubated together for 30 min followed by the addition of 50 μl stop solution with OD recorded at 490 nm by a ELX800 UV microplate reader (Bio-Tek) ^2, 7^. (ii) Inhibition assay for IgE inhibition (determining actual IgE peptide, e.g., nECP): vaccinated PBMCs were harvested on day 7 to day 10 and mixed with 1x10^4^ SKO7 (typed: HLA-A: 02.01.01G, 03.01.01G; HLA-B: 07.02.01G, 40.01.01G), or EBV JY* (HLA-A2.01, HLA-B7.02) at a 10:1 to 20: 1 E/T ratio and the supernatants were harvested at different time points for analyses in a human IgE ELISA, and human IgG (BioLegend, Inc.).

### II. MoDCs for Immunogenic vs. Tolerogenic DCs

#### 1 Preparation of MoDCs

##### Materials

HLA-A2.01+ and/or HLA-B7.02+ PBMCs were obtained from the San Diego Blood Bank, gender, ethnically balanced, blinded, freezing media (10% DMSO in FBS), complete RPMI1640 media (containing 10% FBS, 2 mM L-glutamine), GM-CSF (1:40 aliquot at 20,000 U/µl; used at 100 U/ml), IL-4 (1:10 aliquot at 600 U/µl; used at 200 U/ml), IL-4, PGE-2 (stock at 2.5 μg/ml), IL-1ß (1:10 aliquot at 20 µg/ml), TNF-α (1:10 aliquot at 20 µg/ml).

##### 1. Regular MoDCs

PBMCs from 1 ml frozen vial were thawed by slow addition of fresh DMEM media .1, .2, .4, .8, 1, 10 ml media with 1 min between, centrifuged at 700 rpm for 5 min. The pellets are resuspended in 5 ml complete RPMI 1640 media and added a 10 cm dish for 4 h. The nonadherent cells are decanted. The adherent fractions are incubated with 100 U/ml GM-CSF and 200 U/ml IL-4 for five days for immature MoDCs, and on day 6, maturation cocktails: 1 µg/ml PGE-2, 5 ng/ml IL-1ß, and 10 ng/ml TNF-α are added for overnight incubation for mature MoDCs. At the end of the maturation period (day 3), cells were seeded in a 96-well flat bottom plate (1x10^5^/well), pulsed with SP-1/SP-2, and/or A2-32 (VTLGCLATGY at 10 µg/ml) and measles helper peptide (10 μg/ml) (ISISEIKGVIVHKIETILF) for 8 h, and then mixed with NAD 3 X10^5^ (FastDCs: NAD = 1:3). The nonadherent decanted cells are centrifuged and resuspended in 1 ml freezing media, stored at -80°C as sources of NAD.

##### 2. FastDCs

Fast DCs were prepared by culturing the afore-mentioned adherent PBMC in complete RPMI 1640 medium (containing 10% FBS, 2 mM L-glutamine) with 100 units/ml GM-CSF and 200 units/ml IL-4 for 24 h to obtain immature FastDCs. Maturation was induced for an additional 48 h incubation in the presence of a maturation cocktail containing IL-1ϕ3 (5 ng/ml), IL-6 (60 ng/ml), TNF-α (10 ng/ml), and PGE2 (1 µg/ml). At the end of the maturation period (day 3), cells were seeded in a 96-well flat bottom plate (1x10^5^/well), and stimulated with nECP peptides and helper peptides as described above. and then mixed with NAD 3 X10^5^ (FastDCs: NAD = 1:3).

#### 2 FACS Cell Staining Procedures

##### Extracellular staining

###### Materials

U or V bottom 96 well plate, wash buffer (1% BSA in PBS), staining buffer (1% BSA in PBS + 2 µg/ml Trustain FCX), FC blocking reagent (Trustain human/mouse; Biolegend, fluorochromes-labelled staining antibodies, Isotype control antibodies (ex. mIgG1-FITC), Optional: fixation reagent (2% paraformaldehyde).

###### Protocol

(1). Aliquot cells (3x10^5^-2x10^6^) into 96 well plates and spin at 700 RPM for 7 minutes at 4°C. (2). Aspirate the supernatant and wash cells with 200 µl of wash buffer. (3). Aspirate the supernatant and repeat the wash. (4). Aspirate the supernatant and resuspend cells in 40 µl – 2 µl/antibody. (5). Add 2 µl of each staining antibody and mix well. Incubate for 30 minutes at 4°C in the dark. (5a). From this step forward, make sure to minimize light exposure of the cells to protect fluorescent dyes from degrading; or (5b). Prepare compensation beads at this step; add 1 drop of compensation beads with 2 µl of staining antibody and incubate for 30 minutes. (6). Add up to 200 µl with wash buffer and spin for 700 RPM for 7 minutes at 4°C. (7). Aspirate the supernatant and repeat the wash twice. (8). For live cell flow cytometry, resuspend in 200 µl of PBS and move into 5 ml FACS tubes for immediate flow cytometric analysis. (9). Alternatively, resuspend the above washed and pellets from step 7 in 100 µl of PBS and 100 µl of 4% paraformaldehyde, and incubate for 15 minutes in the dark at 4°C. (10). Cells are then washed, twice and resuspend in 200 µl of PBS into a 5 ml tube at 4°C until flow cytometric analysis.

##### Intracellular and nuclear staining

###### Materials

Cells of interest, brefeldin A (used at 5 µg/ml), monensin (used at 2.5 µg/ml), Stimulating reagents, staining reagents (fluorochrome conjugated labelling antibodies, compensation beads (stained with each fluorochrome), 1% BSA in PBS, permeabilization Buffer (1% BSA, 0.1% saponin/Triton X-100, in PBS), dilution buffer (2 µl/ml Trustain FCX, 0.1% saponin/Triton X-100, 1% BSA in PBS), wash buffer (0.1% saponin/Triton X-100, 1% BSA in PBS), and 96 well U-bottom plate.

###### FACS Staining Panel

FACS Staining Panel:
| FACS reagents (BioLegend, Inc.) |  |
| --- | --- |
| Fluorochrome | Biomarkers |
| BV570 | CD4 |
| AF700 | CD8 |
| APC/Cy7 | CD137 (4-1BB) |
| BV421 | TGF-β |
| PerCP/Cy5.5 | CD83 |
| BV605 | CTLA-4 |
| APC | IFN-γ |
| AF488 | FOXP3 |
| PE/Cy7 | IL-10 |
| PE | HLA-B7.02_SP1 Tetramer |
| PE | HLA-B7.02_SP2 Tetramer |

###### Flow cytometric analysis

(1). Add brefeldin A (10 μg/mL) to the FastDC/NAD two stage cultures, or three stage Fast/DC/NAD IgE lineage culture for the final 3 h during incubation. (2). Centrifuge at 300 x *g* for 8 min at 4°C and wash the cell pellets with 200 μL of FACS staining buffer. (3). Resuspend the cell pellet with 100 μL of FACS staining buffer which containing 5 μL Human Trustain FcX and incubate at rt for 5-10 min. (4). Without washing, add 2 μL of fluorochrome-conjugated antibody each for surface marker staining with pipetting to disperse. (5). Incubate mixed cells for at least 30 min at 4°C, protect from light. (6). Wash the cells with 200 μL/well FACS staining buffer for the 96-well plate. Centrifuge at 300 x *g* for 5 min at rt, and discard supernatant; and repeat the step. (7). Add 200 μL of FOXP3 Fixation/Permeabilization working solution to each well and fully resuspended. for 30-60 min at 4°C. (8) Centrifuge samples at 400 x *g* for 5 min at rt, and resuspend the pellets in 200 μL of FACS staining buffer, and repeat permeation/wash two more times. (9) Resuspend pellets finally in 100 μL 1X permeabilization buffer, and incubate with 2 μL of fluorochrome-conjugated antibody each for intracellular for 30 min. (10). Add 200 μL of 1X Permeabilization Buffer to each well and centrifuge samples at 400 x *g* for 5 min at rt and discard the supernatant and repeat this step. (11) Resuspend the cells in 300 μL of FACS staining buffer, analyze by BD LSRII flow cytometer.

### III. Adoptive MoDCs and Immunogenic vs. Tolerogenic Peptide Therapy

Three Stage, Three-Cell Reconstituted Co-Culture for Evaluating IgE Production in Human PBMCs *ex vivo* for Final nECP Vaccine Validation

#### 1 Design Framework and Work Flows

(1). FastDCs are set on day 1, matured on day 2 and modulators added on day 3. (2) Alternatively, modulators are added to day 6 mature MoDCs as a comparative parallel set. (3). NAD will be added at 3 NAD to 1 DC (FastDC or mature MoDC without washing) at 3:1 ratio second stage cocultures. (4). On day 10, 1 DC/3NAD3 will be added to 10 PBMC, and one day prior to mixing to stage 2 cultures, PBMC are stimulated with anti-CD40/IL-4. (4b) Alternatively, the cocultures are added to SKO7 at 1 DC/3NAD: 3 SKO7 ratios. (5). The three stage three cell cocultures are incubated for 7 to 10 days: on day 7, 100 μl supernatants are harvested with 100 μl fresh medium supplementing back; on day 10 all the supernatants are collected for IgE/IgG levels by ELISA. (6) The three stage-three cell cocultures are harvested for FACS analysis

##### Procedure

(1). On day 0, thaw PBMCs (HLA-A2.01-, B7.02+ or HLA-A2.01+/B7.02+multiple donor sources) from liquid nitrogen using hand to warm it up, and transfer to a 15 mL conical tube, carefully diluted the 10% DMSO away by adding 0.1 ml in double increment per minutes until 1 ml addition is reached and quickly add 13 ml RMPI wash medium without FCS. Centrifuge at 700 rpm for 7 min at rt. (2). Remove supernatant completely and resuspend cell pellet with 10 mL of complete RPMI1640 medium with 10% FCS. (3). Plate PBMCs on 10-cm cell culture dish to let the cells attach for 4 h. (4). After 4 h, remove non-adherent (NAD) cells and incubate with fresh RPMI1640 medium at 37°C. (5). Add pre-warmed fresh RPMI1640 medium with IL-4 (200 U/mL, 3.3μl/10 ml) and GM-CSF (100 U/mL, 2μl/10ml) to start differentiation into immature MoDCs (CD45^+^CD83^-^CD14^-/+^) for 1 day. (6). On Day 3, add maturation cocktail with IL-1β (5 ng/mL), TNF-α (10 ng/mL), IL-6 (60 ng/mL) and PGE2 (1 μg/mL) to complete MoDCs maturation process for 2 days. (7). Count cell and seed cell to 96 wells (DC, 1X10^5^, NAD 3X10^5^, 1:3). (8). Treated with SP-1/SP-2 and PADRE or MHP for 8 h to pulse FastDC. (9). Incubated in the presence or absence of various drugs at the optimally titrated concentrations on day 3, 4, 6. (10). On day 7, set up IgE production by IgE-producing cells from 1x10^6^ human PBMCs with 7% H55 human plasma, anti-CD40 antibody (3 µg/mL), IL-4 (300 units/well), IL-6 (60 ng/mL) and IL-21 (60 ng/mL). (11). On day 4 or day 5, thaw SKO7, let them recover at least 3 days. (12). On day 8 or day 10, the stage 2 total cell cocultures will be added to IgE lineage cells (1 DC/NAD: 3 PBMC); (13). Alternatively, the stage 2 cocultures were added to SKO7 at 1 DC/NAD: 1/3 SKO7, and cocultured with IgE lineage cells 7 more days, on day 15 collect 100 μl sup; add back 100 μl fresh media. (14). On day 18 collect all the sup for IgE levels, SP1/SP2. (15). On day 18 left over three-stage cocultures are analyzed by the multi-color FACS. In pilot experiments, non-specific inhibition by natural killer cells (NK), the two stage cocultures are blocked by anti-NKG2D, MICA/MICB and CD94 Abs (10 μg/mL).

### 2 Cell Fractionation

#### Materials

MACS LS Column and MACS αMarker MicroBeads (Miltenyl Biotec), fractionation cell buffer (0.5% BSA, 2 mM EDTA in PBS, pH 7.2), PBMCs.

##### Negative selection

(1). CD4+ T-cells from day 4DC/NAD cocultures were purified using a CD4 isolation kits, human (Miltenyl Biotec; Cat: 130-096-533) negative selection column with magnetic beads coupled with 10 different mAbs, including anti-CD8 with human CD4 T cell untouched, negatively selected in the collected flowthrough. (2). CD8+ T cells were negatively selected using CD8+ T cell isolation kit, human (Miltenyl Biotec; Cat: 130-096-495) with a cocktail of mAbs including anti-CD4. (3). Fractionated CD8+, CD4+, or a CD4+ plus CD8+ of T-cells were then added to the IgE B cell lineage, stimulated with anti-CD40 mAb and cytokines as above, incubated for 7-10 days. (4). supernatants were harvested for human IgE ELISA and IgG ELISA as control.

###### 3 Reconstituted Three-Stage Autologous DCs/Autologous T cells with Autologous Human IgE Producing Cultures

IgE producing cells cultures were initiated with PBMCs obtained from healthy donors selected by the SDBB, Inc. set registry according to self-reporting, wherein the IgE allergy status is not disclosed in the Agreement. PBMC contain multiple B cell lineages, including the IgE lineage and IgG lineage according to immunoglobulin production of the particular isotype, indicating the genetic switch of epsilon heavy chain gene rearrangements ^8, 9^ and commitment of IgE and IgG B cell lineage, e.g., IgE or IgG committed B cells and plasma cells, wherein the capacity of IgE and IgG production were selected in the study. Thus, 1x106 human PBMC (single or preferably double, genotyped HLA-A2.01+/HLA-B7.02+ donors) were established as set up lineage IgE-producing cells using 7% H55 human plasma, anti-CD40 antibody (3 µg/mL) ^10^, IL-4 (300 U/well) ^8, 9^, IL-6 (60 ng/mL) ^11^, and IL-21 (60 ng/mL) ^12^. It is critical to obtain as many human plasmas and routinely screen for the capacity to support consistent human IgE production; with a competent carefully screened and selected plasma, IL-4 is still strictly required for IgE switch, while the cytokine supplementation, IL-6, IL-21 is dispensable. In our hand, all the PBMCs without exception thus far produce high levels of polyclonal IgE (up to 100 ng/ml or more) when cultured with appropriate human plasma. This remains the most important critical observation for enabling the analysis for regulating pathophysiological levels of human IgE production without skewly relying on human IgE producing SKO7 myeloma cells (HLA-A2.01/HLA-B7.02). Performing large batches of human donor plasma screening is rendered possible through the kind help and the generosity of Mr. Howard Brickner, at UCSD, San Diego, CA. The plasma screened for IgE production was treated with heparin with the removal of fibrinogen and clotting factors. This method has produced robust, consistent, and optimal batches for human IgE production for almost all the human PBMCs tested regardless of the diversity of the MHCI haplotypes.

### IV. The Triple Tg Mice HLA-A2.01/HLA-B7.02 in situ DC-based Vaccination

Mice constitutively produced normal levels of chimeric human rodent IgE JW8 in the sera. Transgenic mice were immunized, s.c. with A32/SP-1/SP-2, and MHP known for provide non-cognate helper CD4 in DOTAP liposomes, previously shown for rodent nECP in IgE mice ^13^. For the in vivo assay: Tg mice and littermates (4 each group) were vaccinated the 20 μg each of A32/SP-1/SP-2 with 10 μg MHP in 250 ml, mixed in 250 μl DOTAP liposomes (Roche, Inc.), primed/boost via s.c. at intervals of 10 days. ELISAs for measuring human IgE, was done using an HRP-conjugated anti-human IgE antibody (Bethyl Inc., TX). Rodent IgE was measured using DNP-specific hybridoma IgE as standard, and rat mAb anti-IgE, made by this laboratory ^14^. Spleens, bone marrows (individual mice), mucosal lymph nodes (mediastinal and mesenteric nodes, pooled from 4 mice) prepared day 7 after a booster were mixed at 10:1 E/T ratios with HLA-A2.01+/HLA-B7.02+ SKO7 (ATCC) or HLA-A2.01+/B7.02+EBV JY*A (kindly obtained from Drs. J. Sidney and A. Sette at LJI, La Jolla, CA), and CTL assay performed.

### V. RNA interference to Elucidate Anti-Lipogenic Treg Pathways for DCs-based Treg Homeostasis

### Transfection with siRNA

Immature or mature MoDCs were seeded onto 24-well plate (3 x 10^5^ cells per well in 0.5 ml of 50% DMEM/50% RPMI 1640 media containing 20% FBS) and incubated at 37°C overnight. For each transfection, 2 mixtures were prepared. Mixture A contained 25 μl of the Opti-MEM medium (# 11058-021, Life Technologies) and 1.5 μl of Lipofectamine RNAiMAX (# 13778-030, Invitrogen), and mixture B contained 25 μl of Opti-MEM and 1.5 μl of 10 mM SREBP-1 siRNA (h) (# sc-36557, Santa Cruz Biotechnology) or siRNA-A, a negative control siRNA (# sc-37007, Santa Cruz Biotechnology). The two mixtures were then combined (50 μl), incubated at room temperature for 5 min and then added to cells dropwise. The final concentration of SREBP-1 siRNA or siRNA-A was 30 nM. Transfected cells were incubated at 37°C for 24 hours followed by RNA isolation.

### RNA isolation

Total RNA was isolated using the TRIzol reagent (# 15596026, Life Technologies). In brief, floating cells were pelleted by centrifugation at 700 rpm for 5 min at room temperature in a Beckman-Coulter Allegra 25R centrifuge. To attached cells, add 300 μl of TRIzol per well to lyse the cells. Then transfer the lysate to the corresponding floating cell pellet and mixed well for complete lysis. The lysate was then mixed with 60 μl of chloroform and incubated at room temperature for 2 min followed by centrifugation at 12000 x g for 15 min at 4°C in an Eppendorf 5804 R centrifuge. The aqueous phase was collected and mixed with 150 μl of isopropanol. After incubation at room temperature for 10 min, the mixture was centrifuged at 12000 x g for 10 min at 4°C. RNA pellet was washed twice with 75% ethanol followed by centrifugation at 7500 x g for 5 min at 4°C. After final wash, RNA pellet was air-dried and resuspended in 30 μl of nuclease-free H2O.

### cDNA synthesis

cDNA was synthesized in a 20-ml system containing 4 μl of 5x LunaScript RT SuperMix (# E3010L, New England BioLabs) and 300 ng of total RNA by running the following program in an Eppendorf Mastercycler (Model 5333): 25oC, 2 min; 55oC, 10 min; and 95oC, 1 min). The cDNA was then diluted to 50 μl with nuclease-free H2O.

### qRT-PCR analysis

96-well plates were used for QRT-PCR analyses and each well contained 10 μl of Luna Universal qPCR Master Mix (#M3003L, New England BioLabs), 2 μl of diluted cDNA, 1 μl of 10 mM forward primer, 1 μl of 10 mM reverse primer and 6 μl of nuclease-free H2O. Analysis was performed using a MyIQ Single Color Real-Time PCR Detection System (Bio-Rad) running the following program: 95°C, 5 min; 95°C, 15 sec – 60°C, 30 sec (40x); 95°C, 1 min; 55°C, 1 min and 55°C, 10 sec (80x). The relative fold gene expression was calculated using the 2^-ΔΔ^Ct method.

### Negative control siRNA

siRNA-A (# sc-37007, Santa Cruz Biotechnology).

No sequences available

siRNA for SREBP-1 knockdown:

SREBP-1 siRNA (h) (# sc-36557, Santa Cruz Biotechnology) No sequences available.

qRT-PCR primers:

SREBP-1 (h) Forward: 5’-ACAGTGACTTCCCTGGCCTAT-3’ Reverse: 5’-GCATGGACGGGTACATCTTCAA-3’

FOXP3 (h) Forward: 5′-CACAACATGCGACCCC CTTTCACC-3′ Reverse: 5′-AGGTTGTGGCGGAT GGCGTTCTTC-3′

GAPDH (h) Forward: 5’-TTG GCT ACA GCA ACA GGGTG-3’ Reverse: 5’-GGG GAG ATT CAG TGT GGT GG-3’

### RT-PCR

#### Materials

cDNA templates from purified mRNA (50 µl from 7 samples of MoDC culture), 2X Taq Universal SYBR Green Supermix, Forward Primer (used at 10 µM), Reverse Primer (used at 10 µM), DNase/RNase free water, Thermocycling plate.

#### Protocol

All parts of the protocol must be done in a RNase/DNase free environment. Failure to do so will negatively impact the experiment. 1. Create triplicate 20 µl solutions in the 96 well plate using the table below: 2X Taq Universal SYBR Green Supermix (10.0 µl), Forward Primer (1.0 µl), Reverse Primer (1.0 µl), DNA template (2.0 µl), Nuclease Free Water (6.0 µl) in a total of 20.0 µl 2. Cover the plate with an adhesive cover and spin at 1000 RPPM for 4 minutes at RT. 3. Use the program “qPCR2016.tmo” on the MyIQ RT-PCR detection system as follows:

| Cycle | Repeat | Steps | Ramp Speed | Time | Temperature |
| --- | --- | --- | --- | --- | --- |
| 1 | 1 | 1 | Max | 5 min | 95°C |
| 2 | 40 | 2 | Max | 15s; 30s | 95°C; 60°C |
| 3 | 1 | 1 | Max | 1 min | 95°C |
| 4 | 1 | 1 | Max | 1 min | 55°C |
| 5 | 80 | 1 | Max | 10s | 55°C |

### Western Blot for Evaluating Levels of Lipogenic Gene Products in DCs/MoDCs

### Material

Purified protein or cell lysate, Transblot-SD Semi-Dry Electrophoretic Transfer Cell (# 1703940, Bio-Rad Laboratories, Hercules, CA), Mini-Protean TGX gel (# 4561096, Bio-Rad Laboratories, Hercules, CA), Azure c600 imaging system (Azure Biosystems, Dublin, CA), Protein gel (Bio-Rad Protean TGX mini gel for native gels, Nacalai gels for denaturing gels), Denaturing sample buffer (120 mM Tris HCL pH 6.8, 4% SDS, 02% bromophenol blue, 20% glycerol; add 20 µl of 1 M DTT to 80 µl of 2x sample buffer before running gel), or Native sample buffer (62.5 mM Tris pH 6.8, 40% glycerol, .04% bromophenol blue), CHAPS lysis buffer (50 mM Tris-HCl, pH 6.8, 2 mM EDTA, 1% CHAPS, 10 mM orthovanadate, 20 μg/ml leupeptin, 10 μg/ml pepstatin A, 10 μg/ml aprotinin and 1 mM PMSF; 150 μl per dish), MW marker, gel running materials (gel tank, gel casket, power supply, electrical connection), Immobilon-P PVDF membrane (# PVH00010, MilliporeSigma, Burlington, MA), 10X transfer buffer (glycine 1.92 M, Tris 192 mM; add 20% ethanol for 1X), Blocking buffer (5% BSA or skim milk powder), 10X SDS-PAGE Running Buffer (1.92 M glycine, 250 mM Tris, 1% SDS), or 10X Native Running Buffer (1.92 M glycine, 250 mM Tris), TBST (washing buffer; 20 mM Tris HCL pH 7.5, 137 mM NaCl, .1% Tween-20), roller, primary antibodies, secondary antibodies, avidin-HRP, Western-Ready™ ECL Substrate Plus Kit (# 426316, BioLegend, San Diego, CA).

### Procedures

Human peripheral blood mononuclear cells (PBMCs) were cultured in RPMI 1640 medium (# 25-506, Genesee Scientific, San Diego, CA) containing 10% FBS, IL-4 (# 574004, BioLegend, San Diego, CA; 20 ng/ml) and GM-CSF (# 572903, BioLegend, San Diego, CA; 20 ng/ml) for 5 days (immature MoDCs). For maturation, cells were treated with IL-1β (5 ng/ml), IL-6 (60 ng/ml), TNFα (10 ng/ml) and Prostaglandin E2 (PGE2) (1 μg/ml) for 24 hours (mature MoDCs), VD3 treatment (100 nM Calcitriol, 100 nM Bexarotene or 100 nM SR11237). For drug treatment, 1 ml of immature or mature MoDCs (10^6^) per 3.5-cm dish was incubated with drugs as indicated. After incubation, cells were rinsed with 1x PBS followed by lysis with either 1x SDS sample buffer (150 μl per dish) or CHAPS lysis buffer (50 mM Tris-HCl, pH 6.8, 2 mM EDTA, 1% CHAPS, 10 mM orthovanadate, 20 μg/ml leupeptin, 10 μg/ml pepstatin A, 10 μg/ml aprotinin and 1 mM PMSF; 150 μl per dish). To detect proteins by western blotting, 15 μl of cell lysate (15-30 μg of proteins) per lane were resolved by 4-20% SDS-PAGE at 100 volts for 80 min with a Mini-Protean TGX gel (# 4561096, Bio-Rad Laboratories, Hercules, CA).

Proteins were then transferred to the Immobilon-P PVDF membrane, using Transblot-SD Semi-Dry Electrophoretic Transfer Cell at 15 volts for 45 min. The blot was subjected to incubation with primary antibodies at 4°C overnight followed by washing for 3 x 5 min with TBST. Blots were then incubated with HRP-conjugated secondary antibodies at rt for 1 hour. After 3 x 5 min washing with TBST, blot was incubated with Western-Ready™ ECL Substrate Plus Kit for 5 min at rt. Protein bands were detected and imaged by Azure c600 imaging system. Relative SREBP-1 or SHP levels in gels were quantified by ImageJ (Rasband, W.S., ImageJ, U. S. National Institutes of Health, Bethesda, Maryland, USA, https://imagej.nih.gov/ij/, 1997-2018) with graphs created by Prism 6 (GraphPad Software, San Diego, CA). (a) The SREBP-1 levels: Primary antibody: mouse anti-SREBP-1 monoclonal antibody (# sc-365513, Santa Cruz Biotechnology, Dallas, TX; 1:200). Secondary antibody: HRP-conjugated goat anti-mouse IgG (H+L) secondary antibody (# 32430, Thermo Fisher Scientific, Waltham, MA; 1:1000). (b) The ACC levels: Primary antibody: rabbit anti-ACC monoclonal antibody (# 3676T, Cell Signaling Technology, Danvers, MA; 1:1000). Secondary antibody: HRP-conjugated donkey anti-rabbit IgG (# 406401, BioLegend, San Diego, CA; 1:1000). (c) The SCD-1 levels: Primary antibody: sheep anti-human SCD-1 antibody (# AF7550-SP, R&D Systems. Minneapolis, MN; 1:2000). Secondary antibody: rabbit anti-sheep IgG (H+L)-HRP (#6150-05, Southern Biotech, Birmingham, AL; 1:1000). (d) The FAS/FASN levels: Primary antibody: rabbit anti-FASN antibody (# 3189S, Cell Signaling Technology, Danvers, MA; 1:1000). Secondary antibody: HRP-conjugated donkey anti-rabbit IgG (# 406401, BioLegend, San Diego, CA; 1:1000). (e). The SHP levels: rabbit anti-NR0B2 (SHP) polyclonal antibody (# ab96605, Abcam, Waltham, MA; 1:1000), Second antibody: HRP-conjugated donkey anti-rabbit IgG (# 406401, BioLegend, San Diego, CA; 1:2000). (f) The GAPDH levels, HRP-conjugated rat anti-GAPDH antibody (# 607904, BioLegend, San Diego, CA; 1:1000).

#### ELISA for Inflammatory Biomarkers

### Materials

96-well plate (Thermo Scientific Nunc MaxiSorp Surface; Cat # 469949), Coating buffer (15 mM Na2CO3, 35 mM NaHCO3, 0.03% NaN3), Coating antibody: Purified anti-human IgE antibody (BioLegend; Cat # 325502; Clone: MHE-18; 0.5 mg/mL), Standard: Human IgE DES (1 mg/mL), Samples (culture supernatants), Blocking buffer (5% FBS, 1% BSA, 0.1% NaN3, 0.05% Tween-20 in PBS), Washing buffer (PBS with 0.05% Tween-20), ELISA Diluent: 10-fold dilution from Blocking buffer with PBS (0.5% FBS, 0.1% BSA, 0.01% NaN3, 0.005% Tween-20 in PBS), Detection antibody, Goat anti-Human IgE Antibody HRP-Conjugated (Bethyl Laboratories; Cat # A80-108P; 1 mg/mL), TMB substrate (BioLegend Substrate A + Substrate B= 1:1; Cat # 421101). Procedure: (1). 96-well Nunc MaxiSorp plate was incubated with 100 μL of 2 μg/ml coating antibody at rt for 2 h. (2). The plate was washed, blotted dry and added with 300 μL of blocking buffer at rt for 1 h. 100 μL of 4-fold diluted supernatant samples were added to each well. For standard curves, human IgE DES was serially titrate for final volume of 100 μL in each well at rt for 1 h. (3). Plate was washed, and HRP-conjugated detection antibody was added in 100 μL for each well at rt for 1 h, washed and blotted dry. (4). Finally, 100 μL of complete TMB substrate, mixed 1:1 (v/v) TMB substrate A and B prior to use were added, and read at 630 nm after 15-, 30-, and 45 min incubation.

## Data Availability statement

All data are available in the main text or the supplementary materials.

## Acknowledgements

We gratefully acknowledge the professional assistance of Dr. Xi- Hua Cao, Mr. David Lee, Dr. I-Fan Lin, Mr. Steven Moody, Mr. Theo Schiff, Ms. Yu-Ting Luo, Dr. Wei-Lun Chen, and Dr. Jhang-Sian Yu and the managerial assistance provided by Ms. Lillian Chen and Mr. Khoi Pham.

## Author contributions

Dr. Chen contributes to conceptual foundation and hypothesis, approaches, and experimental executions and team leading; Dr. Zhang contributes to the hypotheses, approaches and experimental executions and team leading.

## Permission to reproduce (for relevant content)

scientific contents are permitted to be reproduced by the scientific community

